# Metabolomics profiling reveals new aspects of dolichol biosynthesis in *Plasmodium falciparum*

**DOI:** 10.1101/698993

**Authors:** Flavia M. Zimbres, Ana Lisa Valenciano, Emilio F. Merino, Anat Florentin, Nicole R. Holderman, Guijuan He, Katarzyna Gawarecka, Karolina Skorupinska-Tudek, Maria L. Fernández-Murga, Ewa Swiezewska, Xiaofeng Wang, Vasant Muralidharan, Maria Belen Cassera

## Abstract

The *cis*-polyisoprenoid lipids namely polyprenols, dolichols and their derivatives are linear polymers of several isoprene units. In eukaryotes, polyprenols and dolichols are synthesized as a mixture of four or more homologues of different length with one or two predominant species with sizes varying among organisms. Polyprenols have been hardly detectable in eukaryotic cells under normal conditions with the exception of plants and sporulating yeast. Our metabolomics studies revealed that *cis*-polyisoprenoids are more prevalent and diverse in the parasite *Plasmodium falciparum* than previously postulated as we uncovered active *de novo* biosynthesis and substantial levels of accumulation of polyprenols and dolichols of 15 to 19 isoprene units. A distinctive polyprenol and dolichol profile both within the intraerythrocytic asexual cycle and between asexual and gametocyte stages was also observed suggesting that *cis*-polyisoprenoid biosynthesis changes throughout parasite’s development. In addition, we confirmed the presence of an active *cis*-prenyltransferase (PfCPT) and that dolichol biosynthesis occurs via reduction of the polyprenol to dolichol by an active polyprenol reductase (PfPPRD) in the malaria parasite. Isotopic labeling and metabolomic analyses of a conditional mutant of PfCPT or PfPPRD suggest that polyprenols may be able to substitute dolichols in their biological functions when dolichol synthesis is impaired in *Plasmodium*.

---

Malaria is caused by protozoan parasites of the genus *Plasmodium* and most cases of lifethreatening malaria are attributable to infection with *Plasmodium falciparum*. The parasite has a complex life cycle that involves the human host and its vector, the *Anopheles* mosquitoes. All clinical features are caused during the asexual intraerythrocytic life cycle due to the repeated invasion of human red blood cells (RBCs). The asexual intraerythrocytic developmental cycle of *P. falciparum* lasts around 48 h, during which the parasite progresses through four morphologically different stages: ring, trophozoite, and schizont stages, ending with rupture of the erythrocyte and release of merozoites that will invade new erythrocytes. Transmission of the malaria parasite requires development of male and female gametocytes (gametocytogenesis), which are ingested by female mosquitoes during a blood meal and undergo sexual reproduction in the mosquito’s midgut. Nondividing *P. falciparum* gametocytes take between 10 and 12 days to fully mature and progress through five morphologically distinct forms (stages I to V), which are different from other *Plasmodium* species.

During their complex life cycle malaria parasites encounter different nutritional environments within and between hosts. The presence of *de novo* and salvage pathways gives parasites a great metabolic flexibility to cope with those changes^1,2^. For example, mature RBCs are capable of only a few metabolic functions since transcription and translation is not present in these cells. However, a wide variety of metabolites are available to the parasite in the human plasma^3^. Among the metabolic pathways that become inactive in mature RBCs is the mevalonate pathway which synthesizes the isoprenoid building blocks isopentenyl diphosphate (IPP) and dimethylallyl diphosphate (DMAPP)^4^ Downstream *de novo cis*-polyisoprenoid biosynthesis of dolichols is also inactive in mature RBCs^5^ but dolichols synthesized during erythropoiesis are still present in erythrocytes^6,7^ On the other hand, *P. falciparum* has active isoprenoid biosynthesis during the asexual intraerythrocytic developmental cycle as well as during gametocytogenesis where the isoprenoid precursors IPP and DMAPP are synthesized through the methylerythritol phosphate (MEP) pathway^8–10^.

The MEP pathway is localized in the apicoplast^11,12^, a unique chloroplast-like organelle essential for growth and pathogenesis of the malaria parasite^13^ (Fig. 1). Moreover, supply of the isoprenoid precursor IPP is the sole metabolic function of the apicoplast in asexual intraerythrocytic cycle and gametocyte stages^10,14^ The fact that exogenous supply of IPP alone allows parasites lacking the apicoplast to normally grow and develop indicates that IPP is transported out of the apicoplast where the synthesis of *cis*-polyisoprenoid products is predicted to occur (Fig. 1)^10,14^ *cis*-Polyisoprenoids (polyprenols, dolichols and their phosphate esters and carboxylic acid derivatives) are linear polymers of several isoprene units. These lipids are present in all membrane systems^15,16^ but their biological functions besides protein glycosylation in the ER remains largely unknown. Recently, *cis*-polyisoprenoids gained special attention due to several breakthroughs in the field including improvements in the analytical techniques for their analysis^17,18^, partial identification of their enzymatic machinery^19^, and the discovery of new biological functions beyond glycosylation^20,21^. Among some of these biological functions are the regulation of the membrane fluidity^21^, stimulation of spore wall formation in yeast^20^, and scavenging free radicals in cell membranes^22–24^. Interestingly, in cancer cells^25^ and *P. falciparum*^26^, protein dolichylation occurs as a post-translational event, but functions of such derivatives and enzymes involved in this posttranslational protein lipid modification remain unknown.

**Figure 1.**
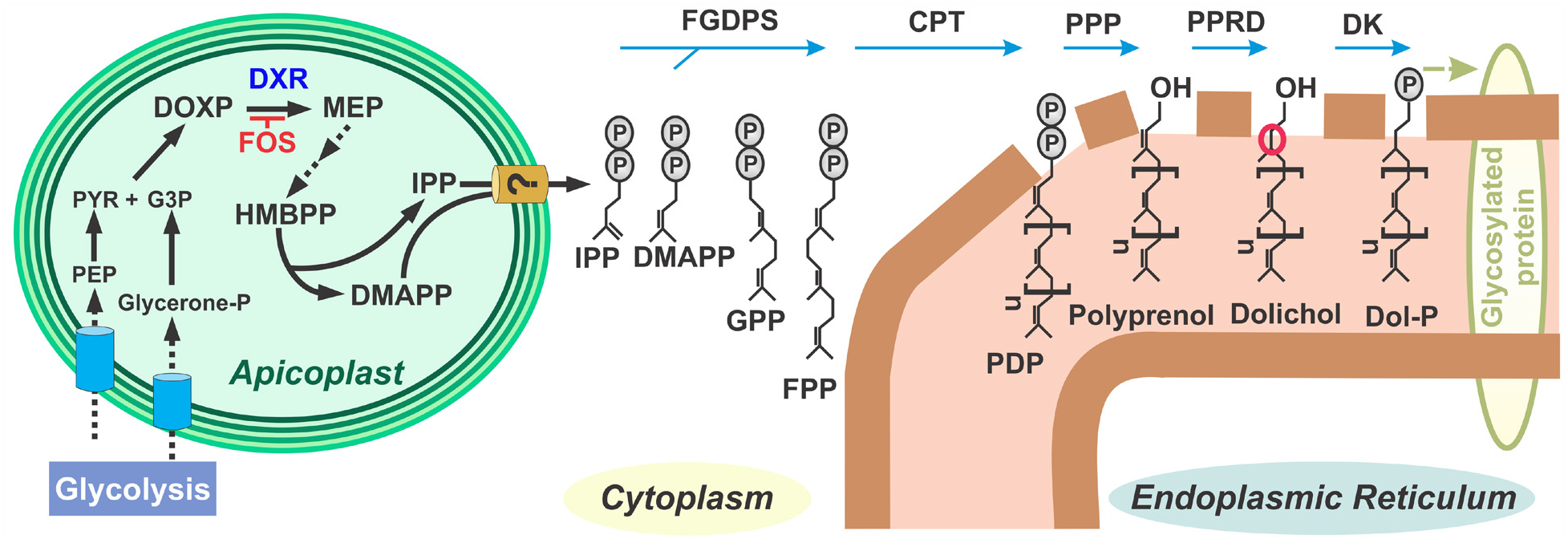
*De novo cis*-polyisoprenoid biosynthesis in *P. falciparum* is predicted to occur in the ER. Phosphoenolpyruvate (PEP); pyruvate (PYR); glyceraldehyde 3-phosphate (G3P); 1-deoxy-D-xylulose-5-phosphate (DOXP); DOXP reductoisomerase (DXR); fosmidomycin (FOS); 2-*C*-methyl-D-erythritol-4-phosphate (MEP); hydroxy-3-methyl-but-2-enyl diphosphate (HMBPP); dimethylallyl diphosphate (DMAPP); isopentenyl diphosphate (IPP); geranyl diphosphate (GPP); farnesyl diphosphate (FPP); farnesyl-geranylgeranyl diphosphate synthase (FGDPS); *cis*-prenyltransferase (CPT); polyprenyl diphosphate (PDP); polyprenyl diphosphate phosphatase (PPP); polyprenol reductase (PPRD); dolichol kinase (DK); dolichyl phosphate (Dol-P). The a-saturated isoprene unit of dolichol is indicated by a red circle.

*cis*-Polyisoprenoid metabolism and its biological functions in *P. falciparum* remain poorly understood. The initial step of *cis*-polyisoprenoid *de novo* biosynthesis in the malaria parasite is common to all isoprenoid products (Fig. 1) where a multifunctional enzyme synthesizes farnesyl diphosphate (FPP) and geranylgeranyl diphosphate (GGPP)^27^. Synthesis of *cis*-polyprenyl diphosphate (PDP) is then proposed to be catalyzed by a *cis*-prenyltransferase (CPT) which adds several IPP molecules in *cis*configuration to FPP or GGPP. This step is called elongation and in animals occurs in the endoplasmic reticulum (ER) where its downstream products dolichyl phosphates (Dol-Ps) serve as lipid oligosaccharide carrier for protein *N*-glycosylation, *C*- and *O*-mannosylation and glycosylphosphatidylinositol (GPI) synthesis (reviewed in^28^). In other organisms, *cis*-polyisoprenoid synthesis also occurs in peroxisomes (rat^29^, yeast^30^), lipid droplets (yeast^20,31^) and chloroplasts (plants, cyanobacteria^21,32^). The postulated final steps of *cis*-polyisoprenoid biosynthesis involve dephosphorylation of PDP to polyprenol followed by reduction of the a-isoprene unit by polyprenol reductase (PPRD), also called SRD5A3 or DFG10 in mammals and yeast, respectively (Fig. 1). Whether dephosphorylation of PDP is catalyzed by one or two enzymes^33^ has not been elucidated in any organism. Moreover, it has also been suggested that an isopentenol or isopentanol, instead of IPP, is added at the last step, which would avoid the need of dephosphorylation or dephosphorylation and reduction, respectively, but it remains to be addressed^15,34^

In eukaryotes, dolichols and polyprenols are synthesized as a mixture of four or more different lengths indicated by the total number of carbons or IPP units, e.g. polyprenol/dolichol 11 or C55, with one or two predominant species (isoprenologues) where the size varies among organisms. Interestingly, polyprenols are hardly detectable in eukaryotic cells under normal conditions with the exception of plants and sporulating yeast. In the malaria parasite, *cis*-polyisoprenoid biosynthesis has received little attention because the relevance of glycosylation in *Plasmodium* was unclear until recently^28^, and a previous report established that the malaria parasite synthesizes dolichols of 11 and 12 isoprene units^35^. However, our preliminary untargeted lipidomic analyses using liquid chromatography combined with high-resolution mass spectrometry (LC-HRMS) system revealed co-occurrence of polyprenols and dolichols which steered us to reexamine *cis*-polyisoprenoid biosynthesis in *P. falciparum*. Further metabolomics and molecular studies revealed that *cis*-polyisoprenoids are more prevalent and diverse than previously postulated. We uncovered active biosynthesis of medium-long polyprenols and dolichols (15 to 19 isoprene units). Moreover, a distinctive profile of polyprenols and dolichols was observed both within the asexual intraerythrocytic cycle and between asexual and gametocyte stages suggesting that their biosynthesis changes throughout parasite’s development. In addition, we confirmed that dolichol biosynthesis occurs via an active PfCPT (PF3D7_0826400) and reduction of the polyprenol to dolichol by PfPPRD (PF3D7_1455900). Two recent functional profiling studies of the *Plasmodium* genome indicate that PfPPRD and PfCPT are predicted to be essential^36,37^. We were able to obtain transgenic strains for inducible knockdown (KD) of PfCPT or PfPPRD but a lethal or severe growth defect phenotype was not observed in both transgenic strains. *De novo* dolichol biosynthesis was still present in the PfCPT knockdown mutants suggesting that some protein expression is still present, thus, explaining the lack of growth defects despite the lower levels of polyprenols and dolichols as compared to controls. However, *de novo* dolichol biosynthesis was undetectable in the PfPPRD knockdown mutant and a pronounced increased in polyprenol levels was observed. This observation together with the lack of a severe growth defect phenotype suggests that polyprenols may be able to substitute dolichols in their biological functions when dolichol synthesis is impaired.

## Results

### Medium-long polyprenols and dolichols are present in the asexual and sexual intraerythrocytic stages of *P. falciparum*

Biosynthesis of dolichol composed of 11 and 12 isoprene units in the schizont stage of malaria parasites has been assessed previously using radiolabeled metabolic precursors followed by thin layer chromatography (TLC) and high-performance liquid chromatography (HPLC) analysis^35^ as well as by purifying unlabeled metabolites by HPLC followed by mass spectrometry (MS) analysis of each fraction^38^. Modulation of dolichol chain length upon a particular physiological condition has been observed earlier. For example, shorter isoprenologues are found in actively dividing cells such as cancer cells compared to not actively dividing cells^39,40^. Thus, we hypothesized that a similar scenario may occur between actively diving asexual stages and nondividing gametocyte stages of *P. falciparum*. To assess this hypothesis, we performed metabolomics analysis by LC-HRMS as we described previously^41^. In the asexual schizont stage, we detected the presence of both polyprenols and dolichols of 15 to 19 isoprene units with isoprenologues 15, 16 and 17 being the predominant species (Fig. 2a) while in stage IV gametocytes dolichols of 15 to 20 isoprene units were also present, but the predominant species were dolichols of 17, 18, 19 isoprene units (Fig. 2b). Under our experimental conditions, polyprenols were almost undetectable in gametocytes. As expected, human RBCs contain low levels of dolichols of 18 to 20 isoprene units with 19 being the predominant isoprenologue (Fig. 2c). This is the first time that the *cis*-polyisoprenoid profile is determined by LC-HRMS in *P. falciparum*, and under our experimental conditions, dolichols and polyprenols of 11 and 12 isoprene units were not detected probably due to their low abundance. To our knowledge, this is also the first time that detectable levels of polyprenols and cooccurrence of polyprenols and dolichols are reported to be present in any other eukaryotic cell besides plants and sporulating yeast. Interestingly, a distinctive temporal profile of polyprenols and dolichols in the asexual intraerythrocytic developmental cycle of *P. falciparum* was also observed. In schizont stages, dolichol/polyprenol ratios were closer to one except for dolichol 15 while ring and trophozoite stages presented considerably higher ratios than schizonts (Table 1).

**Figure 2.**
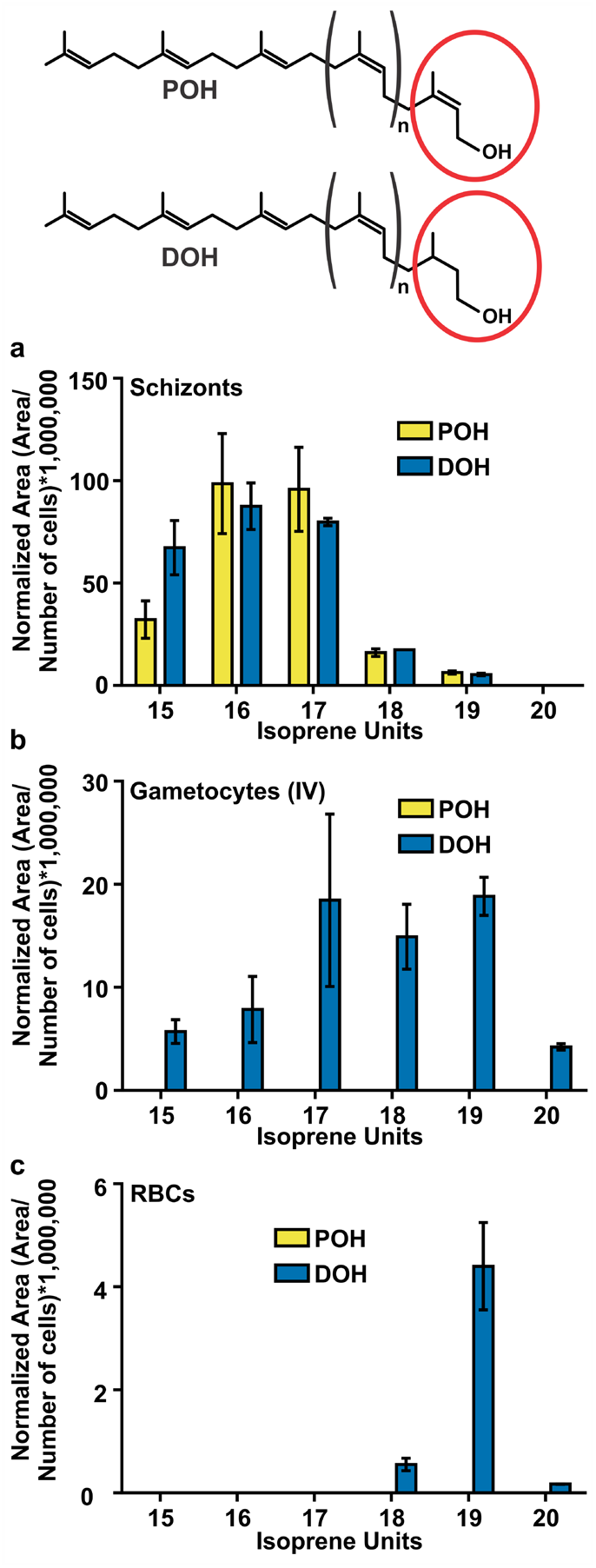
Distribution of polyprenol and dolichol species present in *P. falciparum* schizont stage **(a)**, gametocyte stage IV **(b)**, and uninfected RBCs **(c)** were analyzed by LC-HRMS. Values represent means ± s.e.m. of three independent biological replicates. The structures of polyprenol (POH) and dolichol (DOH) are illustrated where (n) indicates the number of internal *cis*-isoprene units. The α-isoprene unit of polyprenols (unsaturated) and dolichols (saturated) is indicated by a red circle.

**Table 1.** Dolichol to polyprenol ratios (DOH/POH) at each stage of the *P. falciparum* asexual intraerythrocytic cycle. Ratios were calculated using the area of the signal corresponding to each metabolite normalized to the cell number. Values represent the mean ± s.e.m. from three independent biological replicates. POH-ND, polyprenol not detected under the experimental conditions used in this study.

| *P. falciparum*<br>intraerythrocytic stage | DOH/POH-15 | DOH/POH-16 | DOH/POH-17 | DOH/POH-18 | DOH/POH-19 |
| --- | --- | --- | --- | --- | --- |
| Ring | 7.4 ± 2.0 | 9.6 ± 3.1 | 9.2 ± 3.8 | POH-ND | POH-ND |
| Trophozoite | 30.4 ± 6.7 | 36.3 ± 6.1 | 25.6 ± 4.8 | 23.0 ± 3.5 | POH-ND |
| Schizont | 2.1 ± 0.2 | 0.9 ± 0.1 | 0.9 ± 0.2 | 1.1 ± 0.3 | 0.9 ± 0.2 |

### Medium-long polyprenols and dolichols are synthesized *de novo* by *P. falciparum*

In order to confirm that the detected polyprenols and dolichols were in fact synthesized by the parasite, *de novo* biosynthesis was assessed by metabolic labeling using [1-^13^C]glucose or [3-^13^C]IPP (Fig. 3a). The specific labeling pattern of isoprene units resulting from glucose metabolism via glycolysis and incorporation through the mevalonate and MEP pathway are well established^17,18^. Because both the mevalonate and dolichol pathway are inactive in human RBCs^4^, only MEP pathway-specific labeling patterns are observed in *P. falciparum* infected RBCs^4,8,9^.

**Figure 3.**
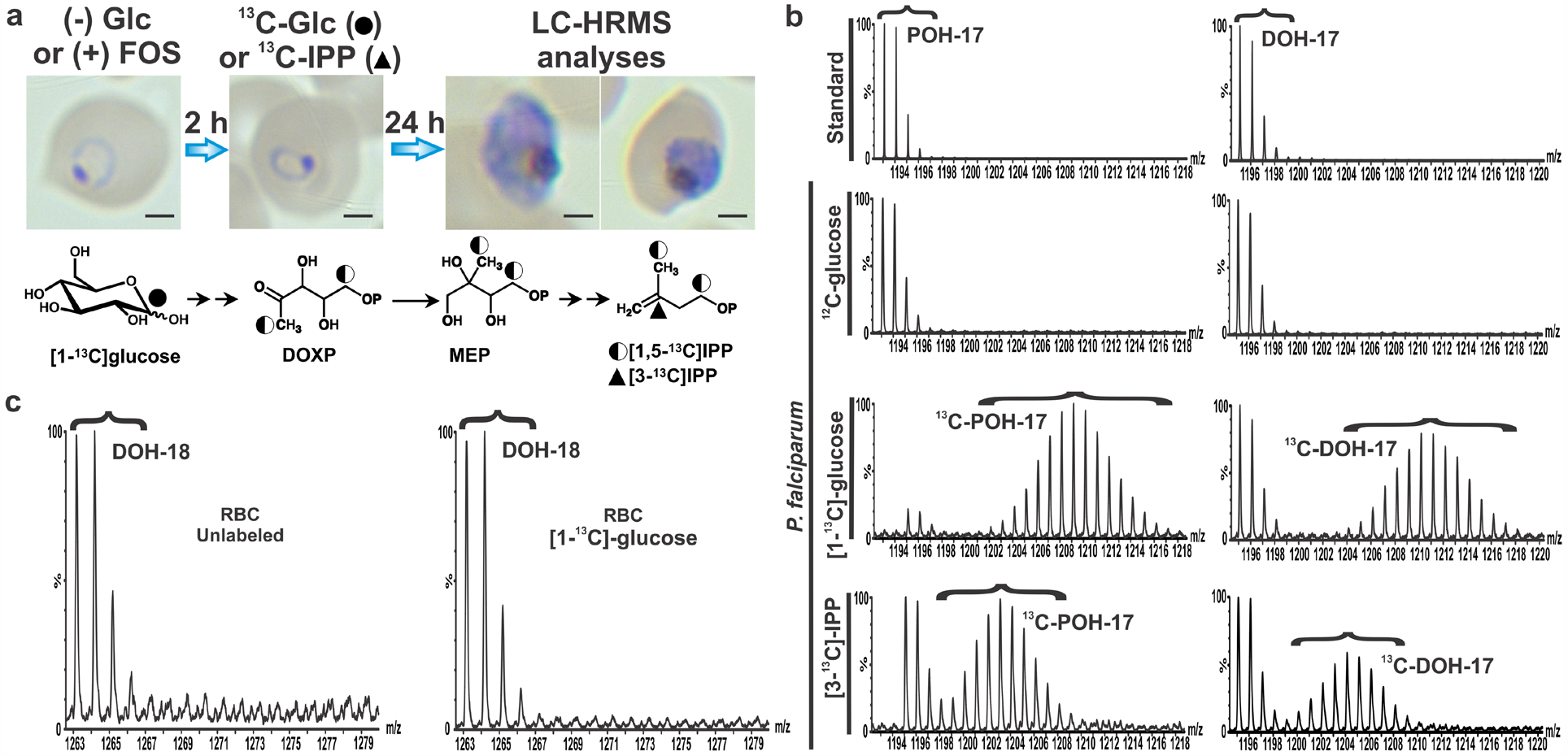
*De novo* biosynthesis of medium-long polyprenols (POH) and dolichols (DOH) in *P. falciparum*. **(a)** Scheme used for metabolic labeling with [1-^13^C]glucose or [3-^13^C]IPP in highly synchronous ring stage cultures. Parasites were recovered at schizont stage for LC-HRMS analysis. A representative Giemsa-stained smear is shown and scale bar indicates 2 μm. The fate of ^13^C through the MEP pathway for [1-^13^C]glucose is showed as a half-black circle to indicate ^13^C abundance, which is 50% of the initial one. A black triangle depicts the localization of the ^13^C atom in the exogenously supplied [3-^13^C]IPP. **(b)** A representative mass spectrum of the standards polyprenol 17 (*m/z* [M+NH4]^+^=1193.1145), dolichol 17 (*m/z* [M+NH4]^+^=1195.1208) and metabolites detected in *P. falciparum* schizont stage is shown in the two upper rows. Distribution of the ^13^C isotopologues observed for native polyisoprenoid alcohols due to the natural abundance of ^13^C is indicated by brackets. Metabolically labeled polyprenol and dolichol are shown in the two lower panels and brackets indicate the *m/z* shift observed as a Gaussian distribution of ^13^C-enriched polyprenol and dolichol isotopologues. **(c)** Mass spectra of dolichol 18 (*m/z* [M+NH_4_]^+^=1263.1868) from uninfected human RBCs cultured in the absence or presence of [1-^13^C]glucose under the same conditions as *P. falciparum* showing lack of *de novo* dolichol biosynthesis.

Highly synchronous infected RBCs in ring stage at 7% parasitemia were first incubated in glucose-free media for 2 h and then supplemented for 24 h with [1-^13^C]glucose (4 g/L) which is known to be metabolized into [1,5-^13^C]IPP (Fig. 3a). A similar scheme was used for the metabolic precursor [3-^13^C]IPP, but in this case cultures were treated for 2 h with 10 μM fosmidomycin (FOS), an inhibitor of 1-deoxy-D-xylulose 5-phosphate reductoisomerase (DXR) – a key enzyme of the MEP pathway (Fig. 1)^8^,^12^, to reduce >75% the endogenous IPP pool as we reported previously^42^, therefore, favoring the uptake of exogenous IPP. Then, cultures were supplemented with a mixture of [3-^13^C]IPP and unlabeled IPP (1:1) at 200 μM final concentration in the presence of FOS. It is well established that 200 μM of exogenous IPP is necessary to reverse growth inhibition caused by FOS or the absence of the apicoplast^10,14,42–44^. During optimization of the ^13^C-biolabeling experimental design, we established that parasites required at least 20-24 h of growth in the presence of the ^13^C-precursor starting at ring stage in order to observe ^13^C-enrichment in polyprenols and dolichols (data not shown). As previously described by Swiezewska and colleagues^17,18^, ^13^C-enrichment in polyprenols and dolichols was observed as a Gaussian distribution of isotopologues for each of the ^13^C-precursors (Fig. 3b, Supplementary Fig. S1). The *m/z* shift toward higher molecular mass mirrors the stochastic distribution of the probability of incorporating differentially labeled ^13^C-IPP into ^13^C-polyprenols, and therefore ^13^C-dolichols, which is determined by the remaining pool of native metabolic precursors in the cell that contain a ^13^C natural abundance of ~1% and the number of ^13^C-atoms incorporated into IPP based on the isotopic dilution due to metabolism^17,45^. As shown in Fig. 3a, the probability of [1-^13^C]glucose to potentially label each of the carbon atoms C-1 and C-5 of IPP is around 0.5, thus, it is expected that the *m/z* shift of the Gaussian distribution will center around [M+(*n*)] with *n* being the number of isoprene units in the polyprenol or dolichol detected as the ammonium adduct. In the case of [3-^13^C]IPP, the *m/z* shift of the Gaussian distribution will center around [M+(1/2 *n*)] of the ammonium adduct because cultures were supplemented with a mixture of [3-^13^C]IPP and IPP (that contains a ^13^C natural abundance of ~1%) at the 1:1 molar ratio to reach the 200 μM final concentration necessary to fully rescue FOS inhibition as previously reported^14^ As expected, only unlabeled dolichols were detected in uninfected RBCs since there is no active *de novo* biosynthesis of isoprenoids (Fig. 3c, Supplementary Fig. S1). Altogether, these results confirmed that the detected polyprenols and dolichols of 15 to 19 isoprene units are exclusively biosynthesized by the malaria parasite.

It has been demonstrated that *P. falciparum* stage III gametocytes incorporate and metabolize ^13^C-glucose^46^. In addition, we previously reported that the MEP pathway intermediates are present in late-stage gametocytes and that IPP synthesis is essential for normal gametocytogenesis since gametocytes lacking the apicoplast require exogenous IPP supplementation until they reach stage IV^10^. Surprisingly, ^13^C-enrichment in dolichols was not detected in stage IV gametocytes when [1-^13^C]glucose was supplemented starting at stage III (Fig. 4). Therefore, these results suggest that synthesis of dolichols occurs earlier than stage III during gametocytogenesis and that IPP is required for other isoprenoid products than dolichols in gametocyte stages III to IV^10^.

**Figure 4.**
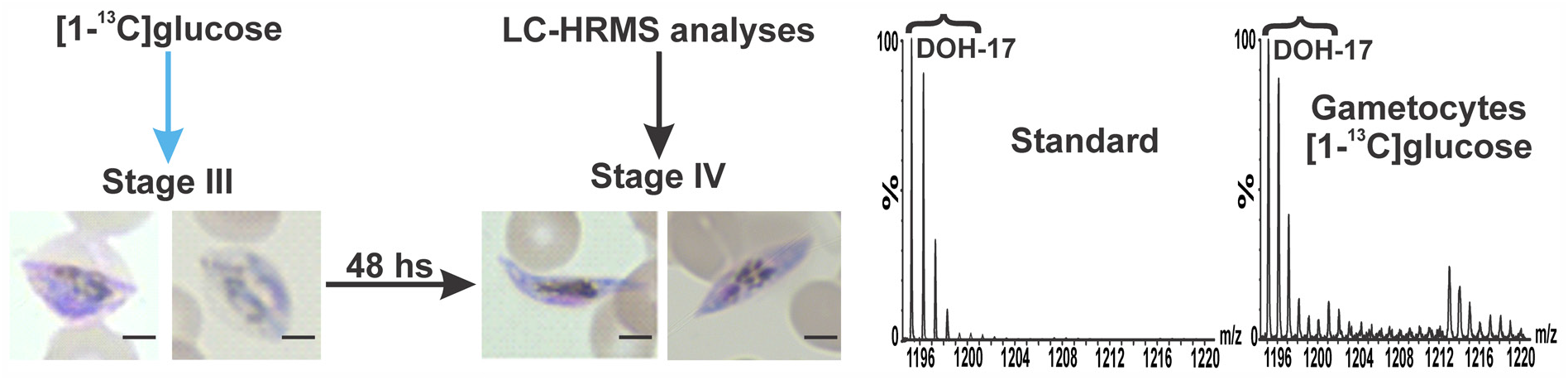
Representative mass spectrum of dolichol 17 (DOH-17) (*m/z* [M+NH_4_]^+^=1195.1208) detected in *P. falciparum* gametocyte cultures metabolically labeled with [1-^13^C]glucose starting at stage III and recovered at stage IV for LC-HRMS analyses as indicated in the scheme. ^13^C-Enrichment was not detected in dolichols from gametocytes under the described experimental conditions. Scale bar indicates 3 μm.

### *P. falciparum* polyprenol reductase restored glycosylation of mCPY and reduced polyprenols levels in yeast *dfg10Δ* mutant cells

We hypothesized that presence of polyprenols in *P. falciparum* indicates that dolichol biosynthesis may occur via reduction of the polyprenol to dolichol by an active PfPPRD (PF3D7_1455900) (Fig. 1). Therefore, the coding sequence of the putative PfPPRD was optimized for codon usage in yeast and its enzymatic function was confirmed by complementation assays in *Saccharomyces cerevisiae*. It was previously reported by Cantagrel and colleagues^47^ that human PPRD (SRD5A3) can complement the *N*-glycosylation defects caused by an inactive yeast DFG10, which is the PPRD in this organism for dolichol biosynthesis. As mentioned above, their downstream products Dol-P serve as lipid linked oligosaccharide carrier for protein glycosylation (Fig. 1). In this work, we used the *dfg10Δ* mutant cells where the *DFG10* gene is deleted from the chromosome and displays a lack of *N*-glycosylation. Mature carboxypeptidase Y (mCPY) has four *N*-glycosylation sites which are all occupied under normal growth conditions in yeast^48^ and depend on a functional PPRD/DFG10 (Fig. 5a)^47^. A single band of fully glycosylated mCPY was observed in wild-type (WT) cells when mCPY is fully glycosylated while four extra bands were detected in *dfg10Δ* mutant cells representing the different glycoforms (−1, −2, −3, −4) (Fig. 5a, lane 2). The mutant transformed with yeast *DGF10* or *PfPPRD* under the control of the endogenous DFG10 promoter showed full correction of the CPY underglycosylation. Some underglycosylated CPY was still detected in cells expressing PfPPRD, suggesting that this enzyme was not as active as yeast DFG10.

**Figure 5.**
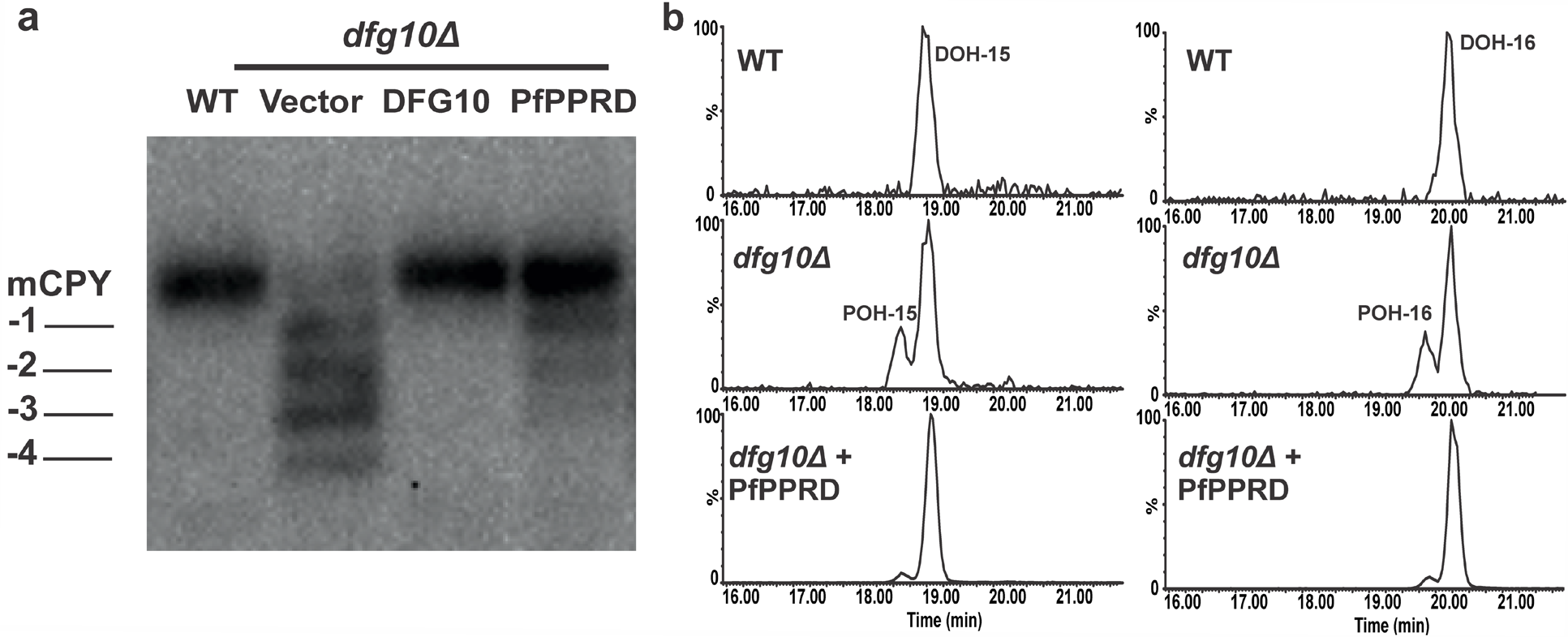
PfPPRD restored *N*-glycosylation of mCPY and reduced levels of polyprenols (POH) in yeast *dfg10Δ* mutant cells. **(a)** Glycosylation status of CPY in *dfg10Δ* mutant cells expressing yeast DFG10 or PfPPRD. Positions of mature CPY (mCPY) and its glycoforms (−1, −2, −3, −4) are indicated. Empty vector (vector) was used as a control. **(b)** Extracted ion chromatogram (EIC) from LC-HRMS analyses of yeast strains showing that for WT only dolichol (DOH) of 15 and 16 isoprene units are detected while in *dfg10Δ* mutant strain dolichols are accompanied by their respective polyprenols 15 and 16. Yeast *dfg10Δ* mutant strain transformed with PfPPRD showed considerable reduction of polyprenols 15 and 16.

In order to confirm the biochemical effects underlying the yeast complementation, we assessed the profile of polyprenols and dolichols by LC-HRMS (Fig. 5b). As described previously^5,47^, only dolichol 15 and 16 were detected in WT yeast while polyprenols along with dolichols were detected in the *dfg10Δ* mutant cells. As expected, *dfg10Δ* mutant cells expressing PfPPRD showed reduction of the polyprenol levels confirming its predicted enzymatic function (polyprenol → dolichol).

### Modification of the CPT and PPRD locus in *P. falciparum*

Two recent functional profiling studies of the *Plasmodium* genome indicate that *Plasmodium* PPRD and CPT are predicted to be essential^36,37^. Thus, in order to gain insight into the polyprenol and dolichol biosynthesis in *P. falciparum*, we used the CRISPR/Cas9 with the tetracycline repressor (TetR) and development of zygote inhibited (DOZI) fusion protein to generate a TetR-DOZI inducible knockdown (KD) mutant strain^49^ of PfCPT or PfPPRD (Fig. 6a). We successfully modified the PfPPRD genomic locus of 3D7 strain (Fig. 6b). We attempted to generate a *C*-terminus triple-human influenza hemagglutinin (HA) tagged PfPPRD to assess its localization but multiple attempts with this construct failed (Supplementary Fig. S2) suggesting that PfPPRD tagged at the *C*-terminus is not efficiently translated or the protein is unstable. It is also possible that the highly conserved catalytic domain of PfPPRD is located in the *C*-terminal as predicted for the human and plant orthologs^47,50^ and tagging interferes with proper enzymatic function. On the other hand, we were able to generate a *C*-terminus triple-HA tagged PfCPT in 3D7 strain (Fig. 6b). Both *P. falciparum* transgenic strains were subjected to clonal selection and genomic modifications were confirmed by DNA sequencing.

**Figure 6.**
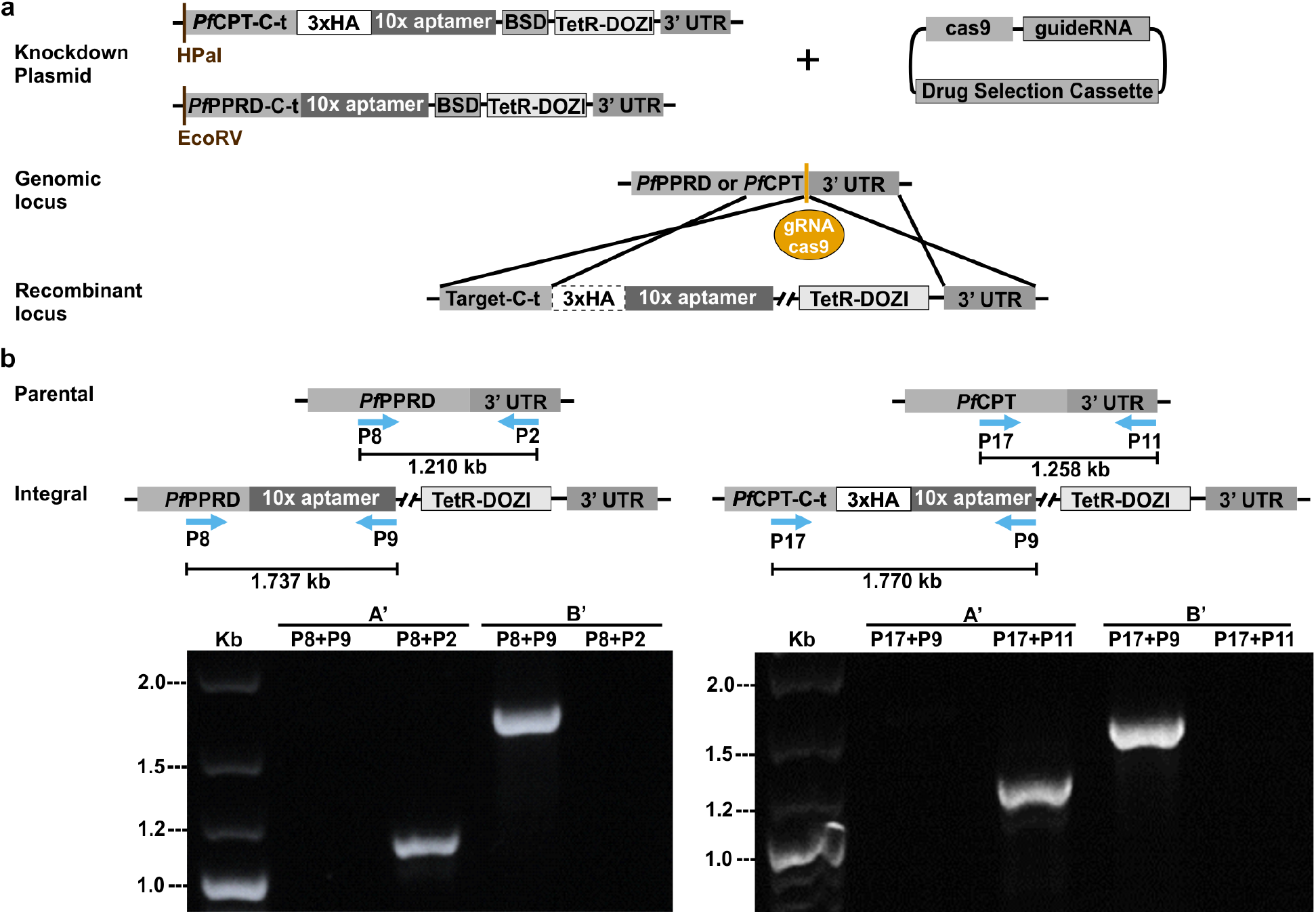
Successful generation of a *P. falciparum* PfPPRD-TetR-DOZI or PfCPT-HA-TetR-DOZI conditional knockdown. **(a)** Schematic representation of plasmids used and integration of the linearized TetR-DOZI plasmid with a 10x aptamer system into *PfPPRD* and *PfCPT* genes. The endonuclease cas9 promotes a double-stranded break in the genomic locus while the repair plasmid provides a homology region at the *C*-terminus of *PfPPRD* or *PfCPT* (target-C-t) for double crossover homologous recombination to introduce a 3’-untranslated region (UTR) RNA aptamer sequence that binds a tetracycline repressor (TetR) and development of zygote inhibited (DOZI) fusion protein to generate TetR-DOZI conditional system. **(b)** PCR analysis confirmed integration at the recombinant locus. Wild type parasites were detected using primers P8-P2 for PfPPRD and P17-P11 for PfCPT (see Supplementary Table S1) after 4 days of transfection (A’). Conditional knockdown parasites were detected after 30 days post-transfection (B’) using primers P8-P9 for PfPPRD and P17-P9 for PfCPT.

### Characterization of PfPPRD and PfCPT conditional knockdowns

The untagged conditional KD system of PfPPRD (PfPPRD-TetR-DOZI) was induced by removing anhydrotetracycline (aTc), which relieves translational repression by the TetR-DOZI complex^49^ (Fig. 7a). Interestingly, morphological defects and a slight but significant growth defect were observed only after 8 days (5^th^ intraerythrocytic life cycle) without aTc (Fig. 7b). We were not able to obtain a PfPPRD-HA-TetR-DOZI strain (Supplementary Fig. S2), thus, it was not possible to determine whether the observed growth and morphological defects were attributed to a gradual or incomplete reduction in PfPPRD protein levels as this enzyme is predicted to be essential^37^. However, the levels of polyprenols and dolichols were determined by LC-HRMS to assess whether PfPPRD enzymatic activity was reduced in the absence of aTc confirming that PfPPRD synthesizes dolichols *in vivo*. Metabolomics analyses were performed in trophozoite stage parasites where the highest dolichol/polyprenol ratios are observed due to low levels of detectable polyprenols (Table 1). In the absence of aTc for 3 days, parasites displayed a pronounced accumulation of polyprenols 15 to 18 and a slight reduction of dolichols 15 to 17 without changes in the levels of dolichol of 18 to 20 isoprene units suggesting that some residual PfPPRD activity may still be present (Fig. 7c). Surprisingly, after 10 days of aTc removal a similar pronounced accumulation of polyprenols 15 to 18 was observed but dolichols 15 to 17 were undetectable while dolichols 18 to 20 were not significantly reduced (Fig. 7d).

**Figure 7.**
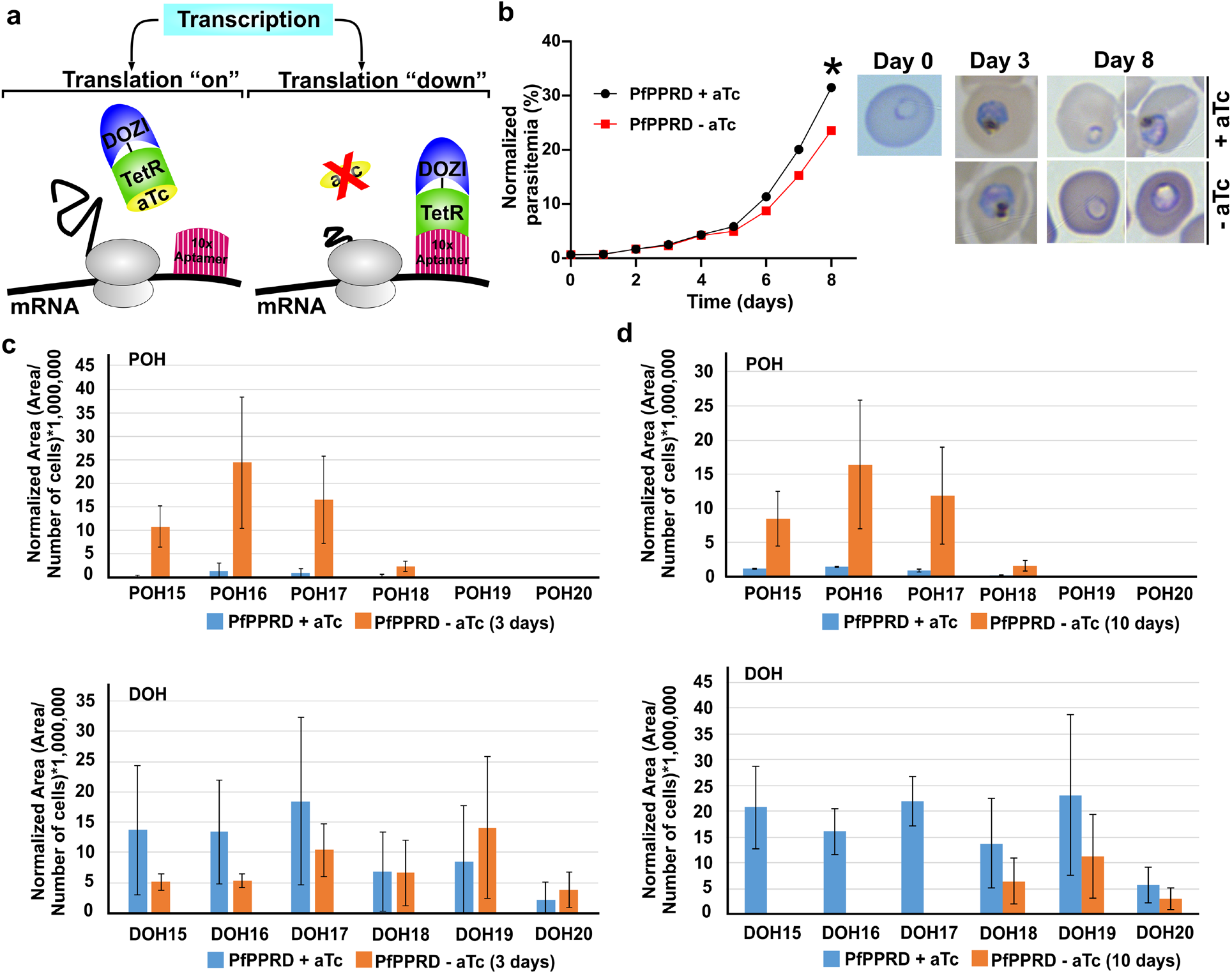
Characterization of PfPPRD conditional knockdown mutant. **(a)** Schematic representation of the PfPPRD conditional knockdown mechanism. The modified *PfPPRD* locus contains a 3 ‘-UTR RNA aptamer sequence that binds to TetR and DOZI fusion protein. In the presence of anhydrotetracycline (aTc), the TetR-DOZI complex is stabilized and the 3’-UTR aptamer is unbound. Removal of aTc causes binding of the TetR+DOZI repressor to the aptamer region which results in reduced protein levels. **(b)** Time-course of growth of PfPPRD-TetR-DOZI knockdown parasites treated with BSD only or BSD+aTc. Data are shown as mean ± SD (n = 3) with error bars smaller than the dots. Individual values are shown in Supplementary Table S2. (*) Indicates *p*<0.0001. A representative Giemsa-stained thin smear of parasites cultured in the presence or absence of aTc showing morphological defects observed in parasites after 8 days of aTc removal. The distribution of polyprenol (POH) and dolichol (DOH) species present in *P. falciparum* PfPPRD-TetR-DOZI were analyzed by LC-HRMS in trophozoite stage after aTc was removed from cultures for **(c)** 3 days and **(d)** 10 days. Values are average ± s.e.m. from three biological replicates.

In order to assess if the detected dolichols of 18 to 20 isoprene units were *de novo* biosynthesized by parasites, we performed metabolic labeling with [1-^13^C]glucose in cultures after 10 days of aTc removal. ^13^C-Enrichment was observed in polyprenol 18 but not in dolichols 18 to 20 suggesting that the detected dolichols in cultures after 10 days of aTc removal were not synthesized *de novo* by parasites (Fig. 8, Supplementary Fig. S3). Consistent with our results shown in Table 1, polyprenol 19 was undetectable in trophozoite stage both with and without aTc supplementation. Dolichols of 18 to 20 isoprene units are present in uninfected RBCs (Fig. 2c) and the treatment with saponin used to isolate whole infected erythrocytes depleted of hemoglobin does not separate the host erythrocyte membrane from the parasite^51^ (see materials and methods). Therefore, we cannot rule out that the detected dolichols of 18 to 20 isoprene units are those present in the erythrocyte membrane, which their detection became more prevalent in the absence of biosynthesis by parasites. Thus, it is possible that the lack of severe growth defect observed in the PfPPRD mutant is due to polyprenols performing the biological functions of dolichols in *P. falciparum*.

To further investigate if lack of polyprenols may be essential for the malaria asexual intraerythrocytic developmental cycle, we performed the same analyses using the conditional KD system of PfCPT-HA-TetR-DOZI. Parasites maintained in the presence of aTc displayed both normal growth and morphology with active *cis*-polyisoprenoid biosynthesis (Fig. 9) supporting that presence of a *C*-terminal triple HA tag does not affect PfCPT function. Removal of aTc for 6 days was not lethal but a striking reduction of polyprenols 15 to 18 and dolichols 15 to 19 was observed confirming the enzymatic function of PfCPT *in vivo* and suggesting that some residual PfCPT activity may still be present in the conditional KD allowing parasites to grow normally as this enzyme is also predicted to be essential^37^. Interestingly, PfCPT-KD (Fig. 9a) phenocopied PfPPRD-KD (Fig. 7d) since dolichols of 18 to 20 isoprene units became the predominant species after several days of aTc removal. We also performed principal component analysis (PCA) where results are displayed as score plots, and each point represents a sample that when clustered together indicates similar metabolite composition based on the metabolomic analysis performed, which in this case was for nonpolar lipids including polyisoprenoids present in the hexane fraction. As expected, a significant change in the parasite metabolome was observed in the absence of aTc (Fig. 9b). As mentioned above, we confirmed the presence of the triple HA tag by DNA sequencing, however, PfCPT-HA could not be detected by western blot, but it was detected by immunofluorescence microscopy (IFA) suggesting low expression (Fig. 9d). Interestingly, strong colocalization of PfCPT-HA with the ER marker PfBiP^52^ was observed in ring stage, but colocalization became very weak in schizont stage suggesting that PfCPT subcellular localization changes with development and that *cis*-polyisoprenoid biosynthesis potentially may occur outside the ER. Expression of PfCPT-HA was faintly detected in parasites after 6 days of aTc removal (Fig. 9d) further supporting our metabolomics analysis suggesting presence of residual PfCPT activity (Fig. 9a).

**Figure 8.**
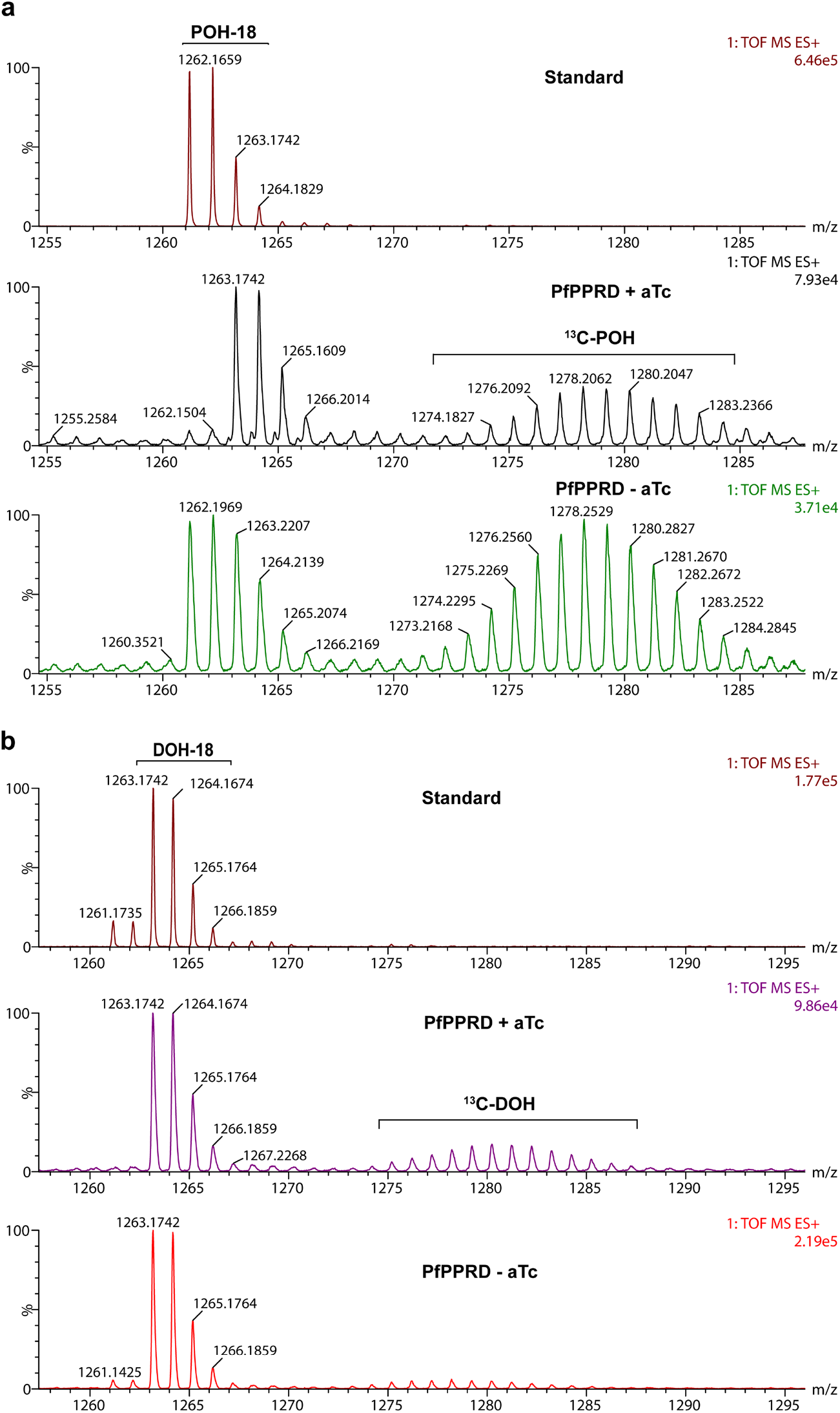
^13^C-Enrichment due to *de novo* biosynthesis in the PfPPRD mutant strain was observed for polyprenol 18 in *P. falciparum* after 10 days aTc was removed **(a)** while ^13^C-enrichment was undetectable for dolichol 18 **(b)** supporting that PfPPRD enzymatic activity is negligible. Metabolic labeling with [1-^13^C]glucose was performed as indicated in Fig. 3a (scheme) and as described in the methodology section. Natural isotopic distribution for polyprenol (POH) and dolichol (DOH) is indicated for the standard. ^13^C-Enrichment from [1-^13^C]glucose is detected as a Gaussian distribution as indicated in the PfPPRD + aTc spectrum. A representative mass spectrum of polyprenol 18 (*m/z* [M+NH4]^+^=1261.1712) and dolichol 18 (*m/z* [M+NH_4_]^+^=1263.1742) is shown.

**Figure 9.**
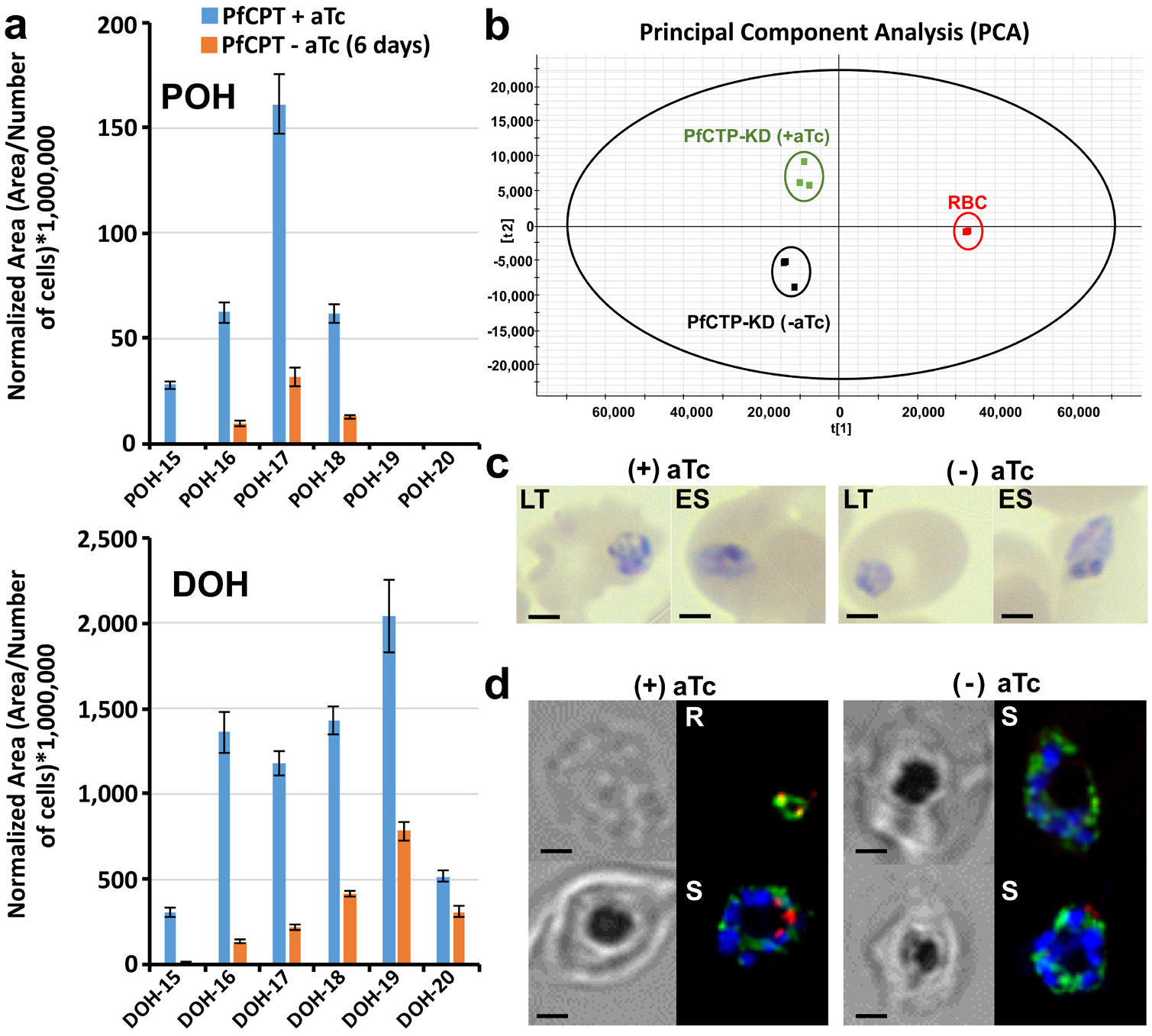
Characterization of PfCPT conditional knockdown mutant. **(a)** Isoprenologues profile in late trophozoite (LT) and early schizont (ES) stage in control (+aTc) and PfCPT knockdown mutant after 6 days of aTc removal. Values are average ± s.e.m. from three independent biological replicates. **(b)** PCA model score plot of LC-HRMS data from three biological triplicates. **(c)** Giemsa-stained thin smear of parasites recovered for LC-HRMS analyses. **(d)** Colocalization with anti-BiP (ER, green), anti-HA (PfCPT, red), and DAPI (nuclei, blue) in asexual intraerythrocytic stages by IFA. R: ring stage, S: schizont stage. Scale bar indicates 2 μm.

### PfCPT and PfPPRD knockdowns have normal MSP1 localization

One of the main known biological functions of dolichols is their role in the synthesis of GPI. The merozoite surface protein 1 (MSP1) is a well-studied GPI-anchored protein uniformly distributed in the parasite plasma membrane and a highly abundant ligand coating the merozoite surface^53^. MSP1 plays a role not only during invasion of RBCs but also during parasite’s egress and it is predicted to be essential^37,53^. Previously, using an inducible Cre recombinase-mediated excision system, it was shown that a truncated MSP1 that lacked a GPI anchor is trafficked to the parasitophorous vacuole but is not attached to the merozoite plasma membrane showing a punctuate localization rather than a uniform distribution^53^. Moreover, these mutants presented a substantially reduced replication rate^53^. Because both PfPPRD-KD and PfCPT-HA-KD presented an altered polyprenol and dolichol profile without a lethal or severe growth defect phenotype, we hypothesized both mutants are able to still carry out protein glycosylation. Therefore, we assessed the localization of MSP1 in PfPPRD-KD and PfCPT-HA-KD by IFA (Fig. 10). As expected, MSP1 showed normal localization as previously reported^53^ in the presence of aTc and no differences were observed in both mutants compared to controls after 10 days of aTc removal suggesting that posttranslational GPI modifications are not affected. Altogether, our results reported here suggest that polyprenols may substitute dolichols as a lipid-linked sugar donor for glycosylation in *P. falciparum* which may explain the lack of a severe growth defect in the PfPPRD mutant and guarantees further investigation.

**Figure 10.**
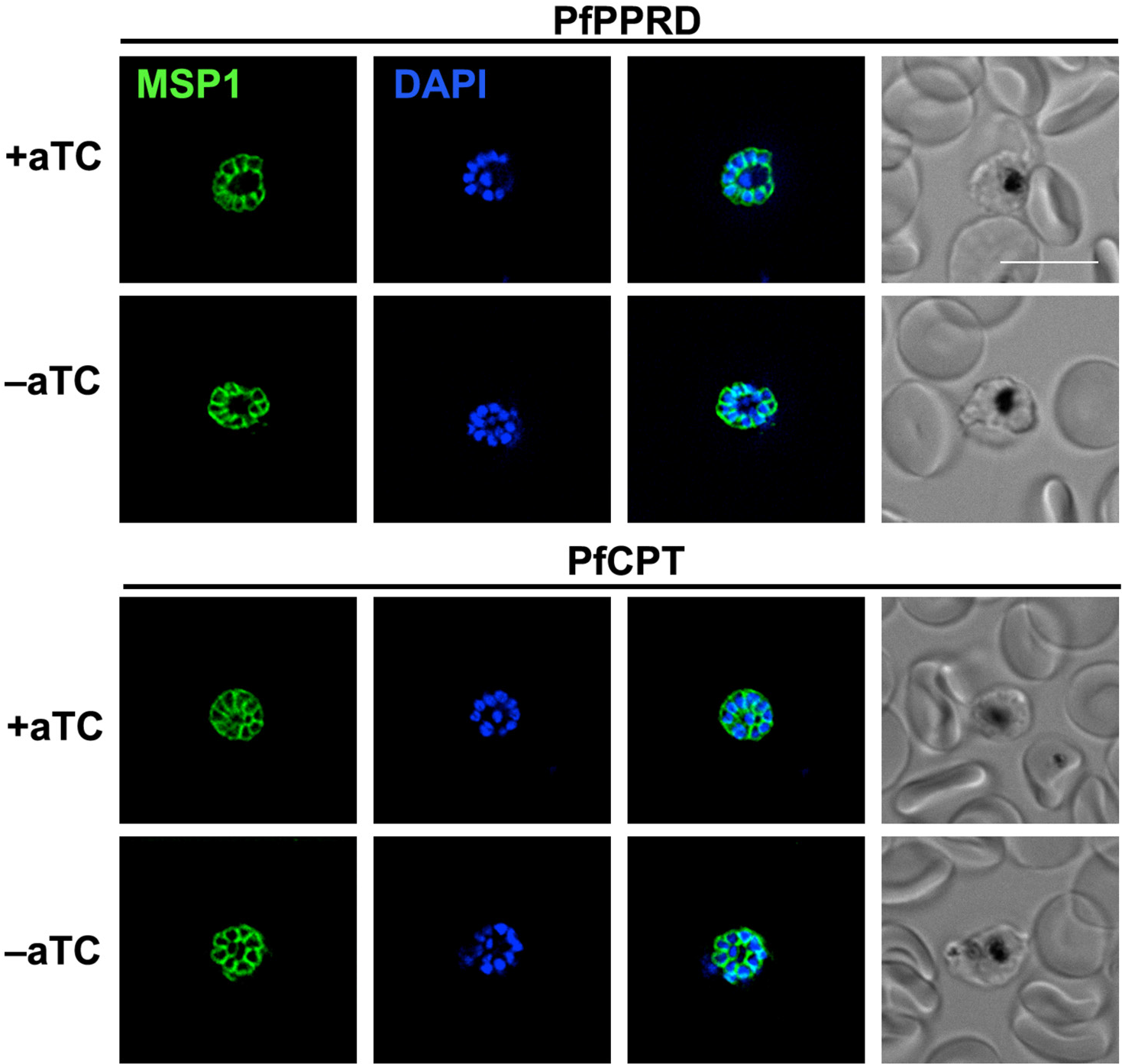
Colocalization of fluorescence signals of anti-MSP1 antibody (green) and DAPI (nuclei, blue) in late schizont stage of PfPPRD or PfCPT mutant strains analyzed by IFA. The aTc supplementation was removed for at least 10 days prior to imagining. Scale bar indicates 10 μm.

## Discussion

Our studies using ^13^C-profiling by LC-HRMS uncovered for the first time in *P. falciparum* cooccurrence of polyprenols and dolichols as well as synthesis of longer dolichol species than previously reported. The presence of a distinctive polyprenol and dolichol profile within the intraerythrocytic asexual cycle and between asexual and gametocyte stages suggest that *cis*-polyisoprenoid biosynthesis changes throughout parasite’s development. Interestingly, in mammals and plants increased level of polyprenol/dolichol ratio results in aberrant glycosylation with shorter glycans^54^. However, it is difficult to assess if a similar response occurs in the malaria parasite since it has been shown that *Plasmodium* synthesizes very short *N*-glycans^28,55^. *cis*-Polyisoprenoid lipids are present in all membrane systems^15,16^ but their biological functions besides protein glycosylation in the ER remain largely unknown. A recent fluorescence anisotropy analysis performed in the protein-dense thylakoid membranes of the chloroplast, revealed that membrane areas with less polyprenols had greater membrane fluidity and photosynthesis was affected^21^. These analyses together with numerous *in vitro* studies employing model membranes^5^ suggest that polyprenols affect the biophysical characteristics of biological membranes. Moreover, it was recently demonstrated in yeast that lipid droplets of vegetative cells contain mainly dolichols of 14 to 17 isoprene units while sporulating cells not only shift toward longer dolichol species (19 to 24 isoprene units) but also accumulate similar levels of polyprenols of 19 to 24 isoprene units. Long-chain polyprenols synthetized by a developmentally regulated CPT present in lipid droplets (Srt1) promote spore wall formation by serving as a signal that activates chitin synthase (directly or indirectly which remains to be determined) possibly through changes in the membrane structure introduced by polyprenols^20^. Therefore, it is possible that the changes in the size of polyisoprene chain and levels of co-occurrence of polyprenols and dolichols during the intraerythrocytic life cycle and gametocyte stages may be associated with their participation in other cellular processes beside glycosylation which guarantees further investigation.

The presence of polyprenols in *P. falciparum* supports that dolichol biosynthesis occurs via reduction of the polyprenol to dolichol and we confirmed that this step is catalyzed by PfPPRD (PF3D7_1455900) which is predicted to be essential in blood stages^37^. Intriguingly, our PfPPRD mutant strain revealed that after aTc has been removed for several days, shorter dolichols were undetectable while dolichols of 18 to 19 isoprene units were still present but not synthesized *de novo*. Due to limitations of current tools to study *cis*-polyisoprenoids metabolism and their biological functions as well as in the approaches to separate parasites from its host erythrocyte membrane, it cannot be rule out that the detected dolichols of 18 to 20 isoprene units are in fact present in the erythrocyte membrane and are not internalized by the parasite. Interestingly, PfPPRD knockdown mutant was not lethal but after at least 8 days of aTc removal a slight but significant growth defect as well as abnormal parasites were observed. Metabolomics analysis revealed that polyprenols were drastically increased in the PfPPRD knockdown mutants. As mentioned above, numerous *in vitro* studies employing model membranes^5^ suggest that polyprenols affect the biophysical characteristics of biological membranes. Therefore, it is also possible that the high levels of polyprenols detected in the absence of aTc may have affected membranes and their functions, thus, contributing to the slight growth and morphological defects observed. In addition, it cannot be ruled out that polyprenols may be able to perform some of the functions of dolichols when their synthesis is compromised including glycosylation as it has been previously suggested to occur in fibroblasts from patients carrying a homozygous SRD5A3 mutation, the human orthologue of PfPPRD^54^ as well as in Chinese hamster ovary cells^56,57^. The efficiency of glycosylation by polyprenols in mammalian cells seems to be lower compared to dolichols and detrimental for cells^54^. Therefore, it is possible that a similar scenario is occurring in the PfPPRD mutants which may also contribute to the slight growth defect observed. However, we did not observed differences in the localization of MSP1 in the PfCPT and PfPPRD knockdown strains when compared to controls suggesting that GPI biosynthesis is not affected in these mutants. As mentioned above, MSP1 lacking a GPI anchor is trafficked to the parasitophorous vacuole but is not attached to the merozoite plasma membrane showing a punctuate localization rather than uniform distribution as we observed in our studies^53^. In the case of the PfPPRD mutant, normal localization of MSP1 suggests that polyprenols may be able to carry out glycosylation in the absence of dolichols (Fig. 7 and 10), if the dolichols of 18 to 19 isoprene units detected in the absence of aTc are localized in the erythrocyte membrane and are not internalized by the parasite. On the other hand, normal localization of MSP1 in the PfCPT may be explained by the fact that the knockdown system used was unable to completely eliminate polyprenol biosynthesis suggesting that parasites can carry out normal glycosylation with low levels of dolichols (Fig. 9 and 10). Currently, we are attempting to generate an inducible knockout of PfCPT to further verify its predicted essentiality^37^. Nevertheless, further investigations are needed to verify that polyprenols can be phosphorylated and glycosylated as dolichols in the malaria parasite.

As a result of several breakthroughs in the field of *cis*-polyisoprenoids^17,19,20^, new biological functions of these lipids are starting to be unveiled besides their known role in protein glycosylation. Further studies are still needed to answer why co-occurrence of polyprenols and dolichols is present in the malaria parasite as well as why *cis*-polyisoprenoid species change during the parasite’s life cycle which may reveal new biological functions of these unique lipids.

## Materials and Methods

### Chemicals

The isotopically labeled intermediate [1-^13^C]glucose (99% isotopic abundance) was obtained from Cambridge Isotope Laboratories Inc (Tewksbury, MA, USA). [3-^13^C]IPP (99% isotopic abundance) from Isoprenoids LC (Tampa, FL, USA) was a kind gift from Dr. Dennis Kyle. All chemicals used in the metabolomics studies were LC/MS grade and purchased from Millipore-Sigma (Burlington, MA, USA). RPMI 1640, HEPES, gentamycin and Albumax I were obtained from GIBCO Life Technologies (Thermo Fisher Scientific, Waltham, MA, USA). Glucose, sodium bicarbonate, hypoxanthine (Millipore-Sigma, Burlington, MA, USA). Polyprenols and dolichols mixtures of 13 to 21 isoprene units were obtained from Avanti Polar Lipids (Alabaster, AL, USA). Polyprenols of 11 and 12 isoprene units were obtained from Isoprenoids LC (Tampa, FL, USA). Dolichols of 11 to 14 isoprene units were from the Collection of Polyprenols, Institute of Biochemistry and Biophysics PAS, Warsaw, Poland.

### Parasite cultures

*P. falciparum* 3D7 and NF54 strains were obtained through the MR4 Malaria Reagent Repository (ATCC, Manassas, VA, USA) as part of the BEI Resources Repository, NIAID, NIH. Parasites were maintained at 5% hematocrit in O^+^ human erythrocytes (The Interstate Companies, TN, USA) in RPMI 1640 supplemented with 5 g/L Albumax I, 2 g/L glucose, 2.3 g/L sodium bicarbonate, 370 μM hypoxanthine, 25 mM HEPES, and 20 mg/L gentamycin. Cultures were maintained at 37^°^C under reduced oxygen conditions (5% CO_2_, 5% O_2_, and 90% N_2_).

### Stable isotope labeling of *P. falciparum*-infected and uninfected RBCs for polyisoprenoids profiling

Mycoplasma-free parasite cultures were tightly synchronized by two consecutive treatments with 5% sorbitol (Millipore-Sigma, Burlington, MA, USA). Each biological replicate was obtained from 75 mL of *P. falciparum-infected* RBC cultures at 5% hematocrit and 7% parasitemia. For stage-specific polyisoprenoid profiling, parasites in trophozoite and schizont stages were obtained from *P. falciparum*-infected RBC cultures starting with early ring stage at 7% parasitemia and harvesting parasites at each indicated stage. Parasites at ring stage were obtained from synchronized cultures starting with early rings of *P. falciparum* at 3% parasitemia and recovering parasites after 48 h incubation to avoid collecting cells after sorbitol treatment.

For metabolic labeling experiments using [1-^13^C]glucose, cultures in ring stage (5% hematocrit and 7% parasitemia) were first incubated in glucose-free RPMI 1640 for 2 h and then supplemented with 4 g/L of [1-^13^C]glucose for 22-24 h. RPMI 1640 without glucose was prepared as we described previously^41^. For metabolic labeling experiments using [3-^13^C]IPP, cultures in ring stage were first treated with 10 μM fosmidomycin for 2 h and then supplemented with [3-^13^C]IPP:IPP (1:1, 200 μM) in the presence of fosmidomycin. In all cases, each biological replicate obtained for infected RBCs was accompanied by its corresponding uninfected human RBCs cultures as controls.

In order to assess dolichol and polyprenol profiles in trophozoite stage of the PfPPRD or PfCPT conditional knockdowns, parasites were grown in media supplemented either with 2.5 μg/mL Blasticidin S and 0.5 μM aTc, or only 2.5 μg/mL Blasticidin S for the indicated times (Fig. 7 and 9). ^13^C-Biolabeling experiments were performed under similar conditions to induce the knockdown and then metabolically labeled as described above.

In all cases, infected RBCs were treated with 0.03% saponin in cold phosphate-buffered saline (PBS) solution containing 2 g/L glucose and washed three times with PBS/glucose by centrifuging 7 min at 10,000 *× g* at 4°C. Ten μL were collected in the last wash to count parasites using a Countess cell counter (Thermo Fisher Scientific, Waltham, MA, USA). Cell pellets were flash-frozen and stored at −80°C until metabolite extractions were performed.

Gametocyte stages were obtained using *P. falciparum* NF54 strain grown in human pooled serum (Interstate Blood Bank, TN, USA) to a final concentration of 10%. Briefly, highly synchronous cultures at early ring stage were obtained by sorbitol treatment, purified using a MACS^®^ magnetic affinity column (day 0) and set at 8-12% parasitemia and 5% hematocrit. On day 0 and 1, gametocytogenesis was induced by incubating cultures with 50% spent media and 50% fresh media for 40-48 h, as described previously^58^. After reinvasion, asexual parasites were depleted from culture by addition of fresh media containing 20 U/mL of heparin for 4 days (day 2 to 5) and sorbitol treatment^59^. After day 5, the media was replaced every two days until gametocytes reached stage IV. Development of parasites was monitored by microscopic evaluation of Giemsa-stained thin smears. For metabolic labeling experiments, stage III gametocyte cultures where obtained as described above (day 6 after induction). Then, parasites were incubated in glucose-free media for 2 h before adding [1-^13^C]glucose (4 g/L) and incubated for 24 h when media was changed to add fresh [1-^13^C]glucose (4 g/L) and to complete a total of 48 h labeling until gametocytes reached stage IV.

### Sample preparation and polyisoprenoid profiling by LC-HRMS

Extraction of metabolites and LC-HRMS analysis was performed as we previously described^41^. Briefly, 1 mL cold methanol was added to each sample and three consecutive extractions with 2 mL of hexane followed by pulse-vortex for 1 min and 10 min sonication in an ultrasonic bath. The upper phases were combined, dried under nitrogen and stored at −80°C until LC-HRMS analysis. For LC-HRMS analysis, samples were resuspended in 100 μL of methanol/acetonitrile/2-propanol (60:25:15, v/v) pulse vortex 5 times and sonicate for 10 min. Analysis of polyisoprenoids was performed on an ionKey/MS system composed of an ACQUITY UPLC M-Class, the ionKey source, and an iKey HSS T3, 100 Å, 1.8 μm (particle size), 150 μm × 100 mm column coupled to a SYNAPT G2-Si mass spectrometer (Waters Corporation, Milford, MA, USA). Isoprenoid separation was accomplished as described previously^41^ and a representative extracted ion chromatogram is shown in Supplementary Fig. S4. For unlabeled samples, peak identification, alignment, normalization, and statistical treatment of the data was performed using Progenesis QI software (Nonlinear Dynamics, Waters Corporation, Milford, MA, USA). For ^13^C-biolabeled samples, MassLynx software (Waters Corporation, Milford, MA, USA) was used for data processing and ^13^C-labeling distribution analysis. The area of the analyte (metabolite) peak was normalized to the cell number and the response of each detected metabolite is expressed as the mean and s.e.m. of at least three independent biological replicates.

### Heterologous expression of PfPPRD and analysis of CPY glycosylation in *Saccharomyces cerevisiae* transformants

HA-tagged DFG10 and PfPPRD (optimized for codon usage in yeast) were expressed under the control of the *DFG10* endogenous promoter from a centromeric plasmid, pRS416. Empty vector pRS416 was included as a negative control. *Saccharomyces cerevisiae* strain BY4741 (MATa his3Δ1 leu2Δ met15Δ ura3Δ) and BY4741-based *dfg10Δ* strains were grown at 30°C in synthetic complete (SC) medium containing 2% galactose as the carbon source^60^. Uracil was omitted from the medium to maintain the plasmid. After two passages (36 to 48 h) in SC medium, cells were harvested when the optical density at 600 nm (OD600) reached between 0.4 and 1.0. Two OD600 units of cells were harvest for protein analysis and 25 OD600 units of cells were harvested for analysis of polyisoprenoids by LC-HRMS.

Total proteins were extracted as described previously^61^. Equal amount of proteins was analyzed using a 12% sodium dodecyl sulfate polyacrylamide gel electrophoresis (SDS-PAGE) and transferred to a polyvinylidene difluoride (PVDF) membrane. Mouse anti-CPY (1:5,000; Life Technologies, Thermo Fisher Scientific, Waltham, MA, USA) was used as the primary antibody; horseradish peroxidase (HRP)-conjugated anti-mouse antibody (1:5,000 dilution; Thermo Fisher Scientific, Waltham, MA, USA) together with the Supersignal West Femto substrate (Thermo Fisher Scientific, Waltham, MA, USA) were used to detect the target protein.

### Generation of a *P. falciparum* strain with an inducible knockdown of PfPPRD or PfCPT

Genomic DNA was isolated from *P. falciparum* 3D7 strain and homology regions were PCR amplified using PrimeStar GXL DNA polymerase (Takara Bio Inc., Mountain View, CA, USA). The 3’-untranslated homology region (3’-UTR) was amplified using forward P1 and reverse P2 primers (Supplementary Table S1). The C-terminal homology region was amplified using forward primer P3 and reverse primer with triple-hemagglutinin (HA) tag P4 or without HA tag P5. The 3’-UTR and C-terminal homology regions were combined using forward P1, reverse P4 or P5 primers and PrimeStar GXL DNA polymerase (Takara Bio Inc., Mountain View, CA, USA). The resulted PCR product was cloned into TetR-DOZI plasmid^49^ by sequence- and ligation-independent cloning (SLIC) using T4 DNA polymerase (Thermo Fisher Scientific, Waltham, MA, USA) as previously described^62^. Before transfection, PfPPRD-HA-TetR-DOZI and PfPPRD-TetR-DOZI plasmids were linearized with EcoRV (New England BioLabs, Ipswich, MA, USA). The RNA guide (gRNA) sequence was chosen using ChopChop online platform (http://chopchop.cbu.uib.no/). Primers forward P6 and reverse P7 were annealed and inserted into pUF1-Cas9-guide^63^ as previously described^64^. Briefly, pUF1-Cas9-guide was digested with BtgZI and annealed oligos were inserted using the SLIC method.

The same approach described above was used to generate a conditional knockdown of PfCPT. Briefly, the 3’-UTR region was amplified using primers forward P10 and reverse P11. The C-terminal homology region was amplified using primers forward P12 and reverse with or without HA tag P13 and P14, respectively. After combination of the homology regions, SLIC and cloning, PfCPT-HA-TetR-DOZI and PfCPT-TetR-DOZI products were linearized for transfection using HPaI (New England BioLabs, Ipswich, MA, USA). Primers forward P15 and reverse 16 were used for annealing and insertion of the guide RNA into pUF1 vector previously digested.

Twenty μg of each linearized plasmid was mixed with 60 μg of pUF1-Cas9-guide and precipitated. DNA pellet was dissolved in Cytomix Buffer^49^ and combined with fresh red blood cells (RBCs). The DNA/RBC mixture was electroporated using BioRad Gene Pulser Xcell^™^ system at 0.32 kV and 925 μF. A culture of *P. falciparum* in schizont stage at 10-15% parasitemia was added to the RBCs and parasites were grown in the presence of 0.5 μM aTc (Cayman Chemical, Ann Arbor, MI, USA). Drug pressure was applied 48 hours post-transfection using 2.5 μg/mL Blasticidin S (BSD) (Millipore-Sigma, Burlington, MA, USA). Media was changed daily for the first 7 days and every other day thereafter. *P. falciparum* cultures were split 1:1 every 7 days to provide fresh uninfected RBCs and to perform DNA extraction for integration analysis.

### Growth and morphological assessment of the PfPPRD-TetR-DOZI and PfCPT-HA-TetR-DOZI

In order to assess potential morphological defects due to the reduced expression of PfPPRD and PfCPT, *P. falciparum* cultures treated with BSD only or BSD+aTc as indicated above were monitored by Giemsa-stained smears every 24 h for over 8 days. Potential growth defects were assessed by seeding cultures in a 96 well plate (0.5% early trophozoite stages at 4% hematocrit) and treated with BSD only or BSD+aTc with each condition tested in triplicate. Throughout the course of the experiment, parasites were sub-cultured to maintain the parasitemia between 1-5%. Parasitemia was monitored every 24 hours by flow cytometry using a CytoFLEX S (Beckman Coulter, Hialeah, FL, USA). A sample of 0.5 μL from each well was stained with Hoechst (Thermo Fisher Scientific, Waltham, MA, USA) at 0.8 μg/mL in PBS and data were analyzed using Prism (GraphPad Software, Inc.). Relative parasitemia at each time point was back calculated based on actual parasitemia multiplied by the relevant dilution factors.

### Statistical analysis

The *t*-test and Benjamini and Hochberg procedure were used to analyze differences between different conditions, and a false discovery rate of 0.01 was used to identify statistically significant differences.

### Fluorescence microscopy

Cells were fixed using a mixture of 4% paraformaldehyde and 0.015% glutaraldehyde and permeabilized with 0.1% Triton-X100 as described previously^62^. Primary antibodies used for IFAs in this study were the following: mouse anti-MSP1 antibody (European Malaria Reagent Repository, 1:500), rat anti-PfBiP MRA-1247 (BEI Resources, NIAID, NIH, 1:100) and mouse anti-HA(6E2) (Cell Signaling Technology Inc., 1:100). Anti-mouse and anti-rat antibody conjugated to Alexa Fluor 488 or Alexa Fluor 546 (1:100, Life Technologies, Thermo Fisher Scientific, Waltham, MA, USA) were used as secondary antibodies. Cells were mounted on ProLong Gold with 4’,6’-diamidino-2-phenylindole (DAPI) (Thermo Fisher Scientific, Waltham, MA, USA) and imaged using a Delta-Vision II microscope system with an Olympus IX-71 inverted microscope using a 100x objective A. Image processing, analysis and display were preformed using SoftWorx and Adobe Photoshop. Adjustments to brightness and contrast were made for display purposes.

## Acknowledgements

This work was supported by the National Institutes of Health (AI108819 to M.B.C.) and American Heart Association Postdoctoral Fellowship (18POST34080315 to A.F.). The following reagents were obtained through MR4 as part of the BEI Resources Repository, NIAID, NIH: *P. falciparum*, strain NF54 (MRA-1000), contributed by M. Dowler, Walter Reed Army Institute of research and strain 3D7 (MRA-102) contributed by Daniel J. Carucci. We thank Julie Nelson and the CTEGD core facility for providing access to flow cytometry equipment. We thank Dennis Kyle for the gift of [3-^13^C]IPP. The yeast strains were kindly supplied to E.S. by Vincent Cantagrel. We thank Grant Butschek for comments and corrections.

## Author Contributions

F.M.Z., A.L.V., E.F.M., M.B.C. conceived of and designed the work. F.M.Z., A.L.V., E.F.M., A.F., N.R.H., K.S-T, M.L.F-M performed data acquisition, analysis and validation. F.M.Z., A.F. and V.M. designed and obtain PfCPT and PfPPRD mutant strains. G.H., K.G. and X.W. designed and performed yeast experiments. F.M.Z, A.L.V., E.F.M., X.W., V.M., E.S. and M.B.C wrote the original draft. All authors reviewed and edited the text.

