## Supplemental Material for "Metabolomics profiling reveals new aspects of dolichol biosynthesis in *Plasmodium falciparum*"

From the [1]Department of Biochemistry & Molecular Biology, University of Georgia, Athens GA 30602; [2]Center for Tropical and Emerging Global Diseases (CTEGD), University of Georgia, Athens GA 30602; [3]Department of Cellular Biology, University of Georgia, Athens GA 30602; [4]School of Plant and Environmental Sciences, Virginia Tech, Blacksburg VA 24061; [5]Institute of Biochemistry and Biophysics, Polish Academy of Sciences, Pawinskiego 5A, 02-106 Warsaw, Poland; [6]Laboratory of Experimental Pathology, Health Research Institute Hospital La Fe, Valencia 46026, Spain

[#] Contributed equally to this work
* To whom correspondence should be addressed: Maria Belen Cassera, Department of Biochemistry & Molecular Biology and Center for Tropical and Emerging Global Diseases (CTEGD), University of Georgia, Athens GA 30602;; Tel. (706) 542-5192.

**Keywords:** *Plasmodium,* malaria, polyprenol, dolichol, polyprenol reductase, SRD5A3, LC-HRMS

**Table S1.** Specific primers used to obtain a PfCPT and a PfPPRD conditional knockdowns.

| **Primer label (localization)** | **Primer Sequence** |
|---|---|
| **P1** (3'-UTR -F) | 5'-CTTTCCGGGCGCGCCTTAAGATATATGATAACATTATTTTATATATATATATAATACATTGGTGTAACGTGTTTTTTTAATAAATGCTG-3' |
| **P2** (3'-UTR-R) | 5'-AAATATATTAAAGATATCCTTACATCATGCATTGTTGATTTAATAGGTGATAATTAC-3' |
| **P3** (C-term-F) | 5'-GATATCTTTAATATATTTTTCGTTCTTCTTATTAAACAAAAATATC-3' |
| **P4** (C-term-R-HA) | 5'-ACGTCATAAGGATAGACGTCTCAAGCGTAATCTGGAACATCGTATGGGTAAGCGTAATCTGGAACATCATATGGGTAAGCGTAATCTGGAACATCGTATGGGTACTTAAGCAAAATATATGGGAAGATTATTTTTCTGTTCCTATATTAAG-3' |
| **P5** (C-term-R) | 5'-ACGTCATAAGGATAGACGTCTCACAAAATATATGGGAAGATTATTTTTCTGTTCCTATATTAAG-3' |
| **P6** (guide RNA-F) | 5'-AAGTATATAATATTTAACGTGTTTTTTTAATAAAGTTTTAGAGCTAGAA-3' |
| **P7** (guide RNA-R) | 5'-TTCTAGCTCTAAAACTTTATTAAAAAAACACGTTAAATATTATATACTTA-3' |
| **P8 (**5'-homology region-F) | 5'-TTTGACTCTCATCTTCGATTAGCTAGGCTACGCCCAAAAGG-3' |
| **P9** (aptamer-R) | 5'-GTAGACCCCATTGTGAGTACATAAATATATTATATAAACTAGACTAGG-3' |
| **P10** (3'-UTR -F) | 5'-CTTTCCGGGCGCGCCTTAAGGTGTTATTGTTTTATTTGTTTTTTTTTTGTGTATTTG-3' |
| **P11** (3'-UTR-R) | 5'-TTACTTTTTCCCGTTAACGGGAGCATCAGCTAAAATTAACTTGGGCCCTAATTTTAC-3' |
| **P12** (C-term-F) | 5'-GCTGATGCTCCCGTTAACGGGAAAAAGTAAAATACAATGAAGAGCAATTAGAAATGTAAG-3' |
| **P13** (C-term-R-HA) | 5'-CGTATGGGTACTTAAGGTGTGGACAAGGATTCGAGTATGAAAATATCCAtTTcGTTG-3' |
| **P14** (C-term-R) | 5'-ACGTCATAAGGATAGGTGTGGACAAGGATTCGAGTATGAAAATATCCAtTTcGTTGTTTG-3' |
| **P15** (guide RNA-F) | 5'-TAAGTATATAATATTCCCTTTTTCAAACAACCAAGGTTTTAGAGCTAGAA-3' |
| **P16** (guide RNA-R) | 5'-TTCTAGCTCTAAAACCTTGGTTGTTTGAAAAAGGGAATATTATATACTTA-3' |
| **P17 (**5'-homology region-F) | 5'-ATATTCCCACACAAGAAGAAGAACTAAAATATCCTGATCAT-3' |

**Table S2.** Time-course of growth analyzed by flow cytometry of PfPPRD-TetR-DOZI knockdown mutant strain treated with BSD only or BSD+aTc.

| | **Normalized parasitemia (%)** | |
|---|---|---|
| **Time (day)** | **PfPPRD + aTc** | **PfPPRD - aTc** |
| **0** | 0.65 ± 0.01 | 0.66 ± 0.10 |
| **1** | 0.75 ± 0.09 | 0.71 ± 0.05 |
| **2** | 1.70 ± 0.06 | 1.71 ± 0.06 |
| **3** | 2.51 ± 0.09 | 2.32 ± 0.14 |
| **4** | 4.36 ± 0.10 | 4.19 ± 0.21 |
| **5** | 5.86 ± 0.66 | 4.97 ± 0.62 |
| **6** | 11.35 ± 0.42 | 8.76 ± 0.77 |
| **7** | 20.09 ± 0.48 | 15.25 ± 0.40 |
| **8** | 31.52 ± 0.34 | 23.57 ± 0.41 |

**Fig. S1. *De novo* biosynthesis of medium-long dolichols (DOH) in *P. falciparum*.** A representative mass spectrum of dolichol 15 (*m/z* $[M+NH_4]^+$=1058.9913) and dolichol 16 (*m/z* $[M+NH_4]^+$=1127.0604) is shown. Natural isotopic distribution of dolichol is indicated for the standard. $^{13}$C-Enrichment from each metabolic precursor is detected as a Gaussian distribution indicated on each spectrum ($^{13}$C-DOH).

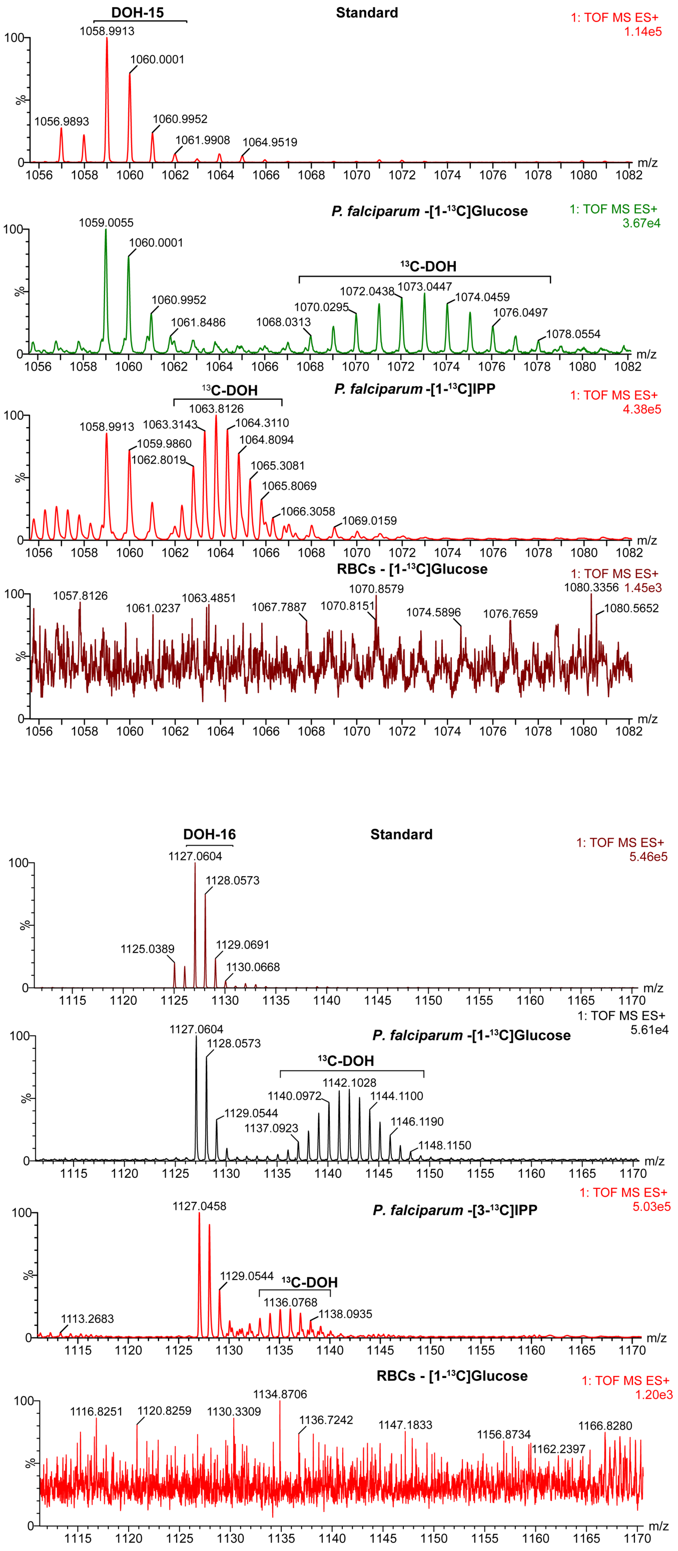

**Fig. S1 (continuation).** A representative mass spectrum of dolichol 18 (*m/z* $[M+NH_4]^+$=1263.1897) and dolichol 19 (*m/z* $[M+NH_4]^+$=1331.2476) is shown.

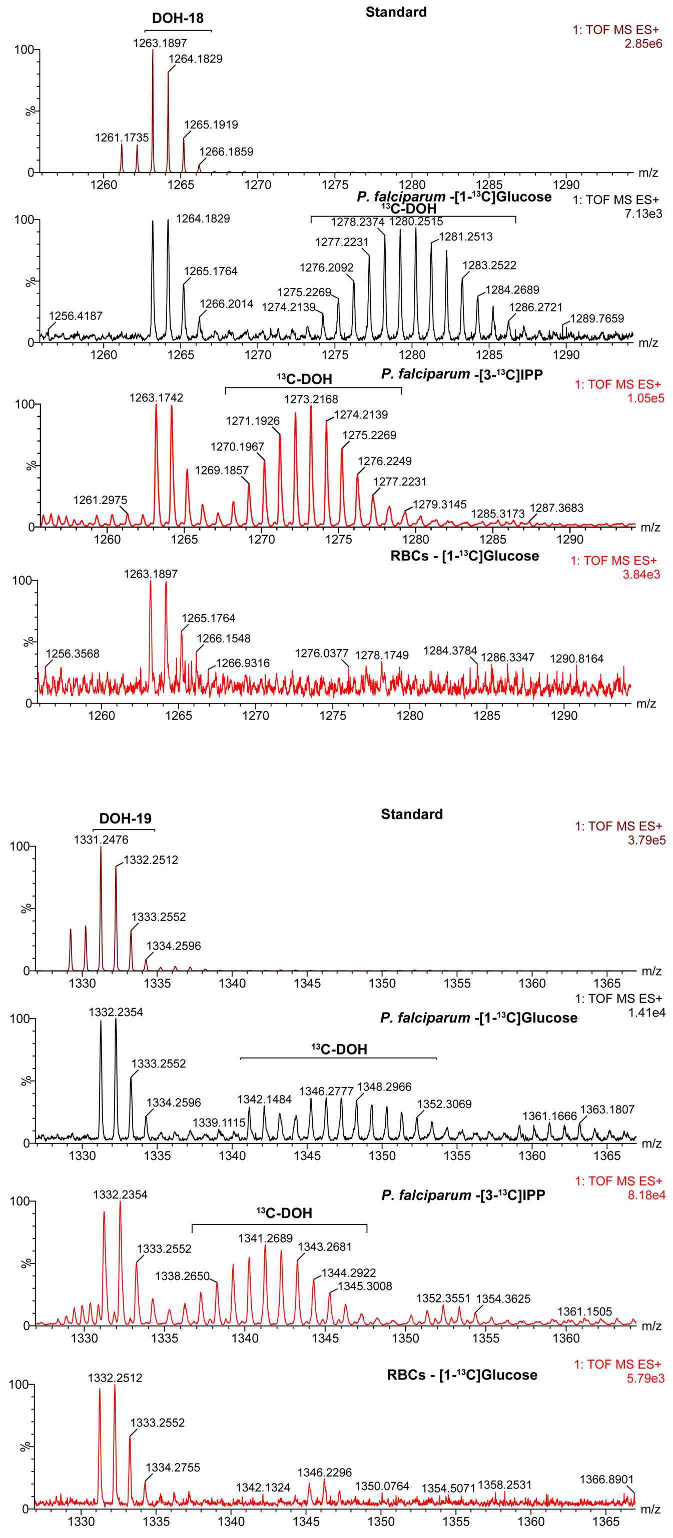

**Fig. S2.** PCR analysis showing that initial integration of the linearized PfPPRD-HA-TetR-DOZI plasmid was observed in the first week of transfection using primers P8 and P9 (see Fig. 6 and Supplementary Table S1), but then the population of parasites harboring PfPPRD-HA-TetR-DOZI was lost in the following weeks suggesting a fitness cost. Wild type parasites were detected using primers P8 and P2 (Supplementary Table S1).

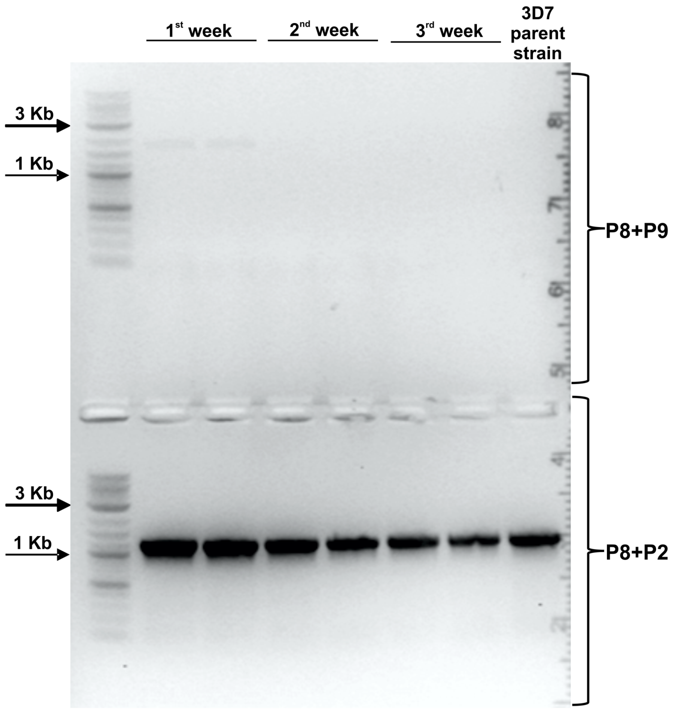

**Fig. S3.** $^{13}$C-Dolichol was not detected in parasites metabolically labeled with [1-$^{13}$C]glucose after 10 days of aTc removal. Metabolic labeling was performed as indicated in Fig. 3a (scheme) and as described in the method section. Natural isotopic distribution for dolichol (DOH) is indicated for the standard. A representative mass spectrum of dolichol 19 (*m/z* $[M+NH_4]^+$=1331.2476) and dolichol 20 (*m/z* $[M+NH_4]^+$=1399.2943) is shown. The area in the spectrum where $^{13}$C-enrichment is expected to appear as a Gaussian distribution is indicated as $^{13}$C-DOH. In *P. falciparum* cultures where aTc was not removed, $^{13}$C-enrichment in dolichol 19 was not detected; however, this result was not unexpected since low levels of $^{13}$C-enrichment using [1-$^{13}$C]glucose were also observed using wild type parasites (see supplementary Fig. S1). A representative Giemsa-stained smear is shown for each condition at the time that parasites were recovered for LC-HRMS analysis.

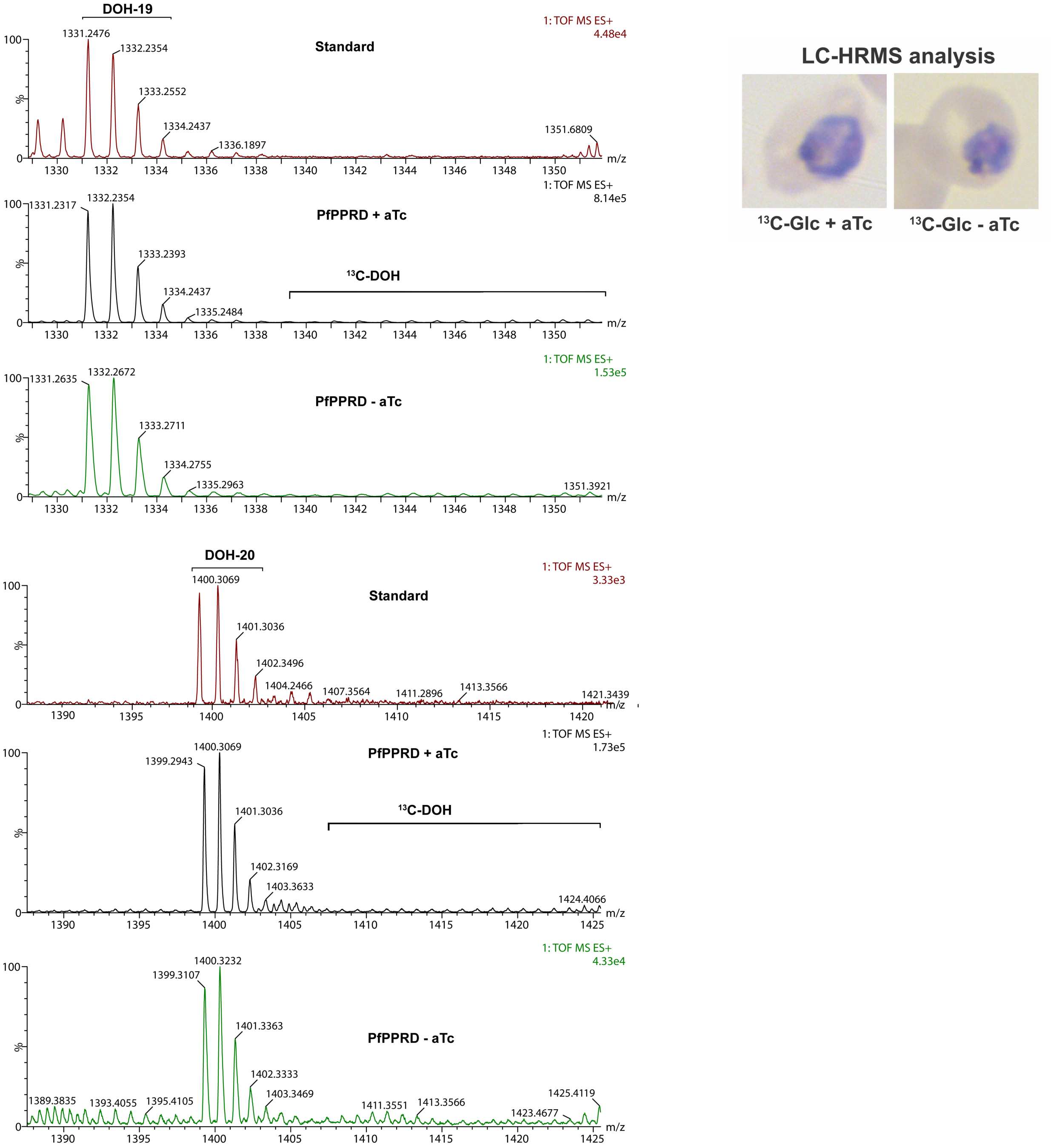

**Fig. S4.** Analysis of polyisoprenoids was performed on an ionKey/MS system composed of an ACQUITY UPLC M-Class and an ionKey source coupled to a SYNAPT G2-Si mass spectrometer. A representative extracted ion chromatogram for polyprenol and dolichol mixtures from Avanti Polar Lipids is shown.

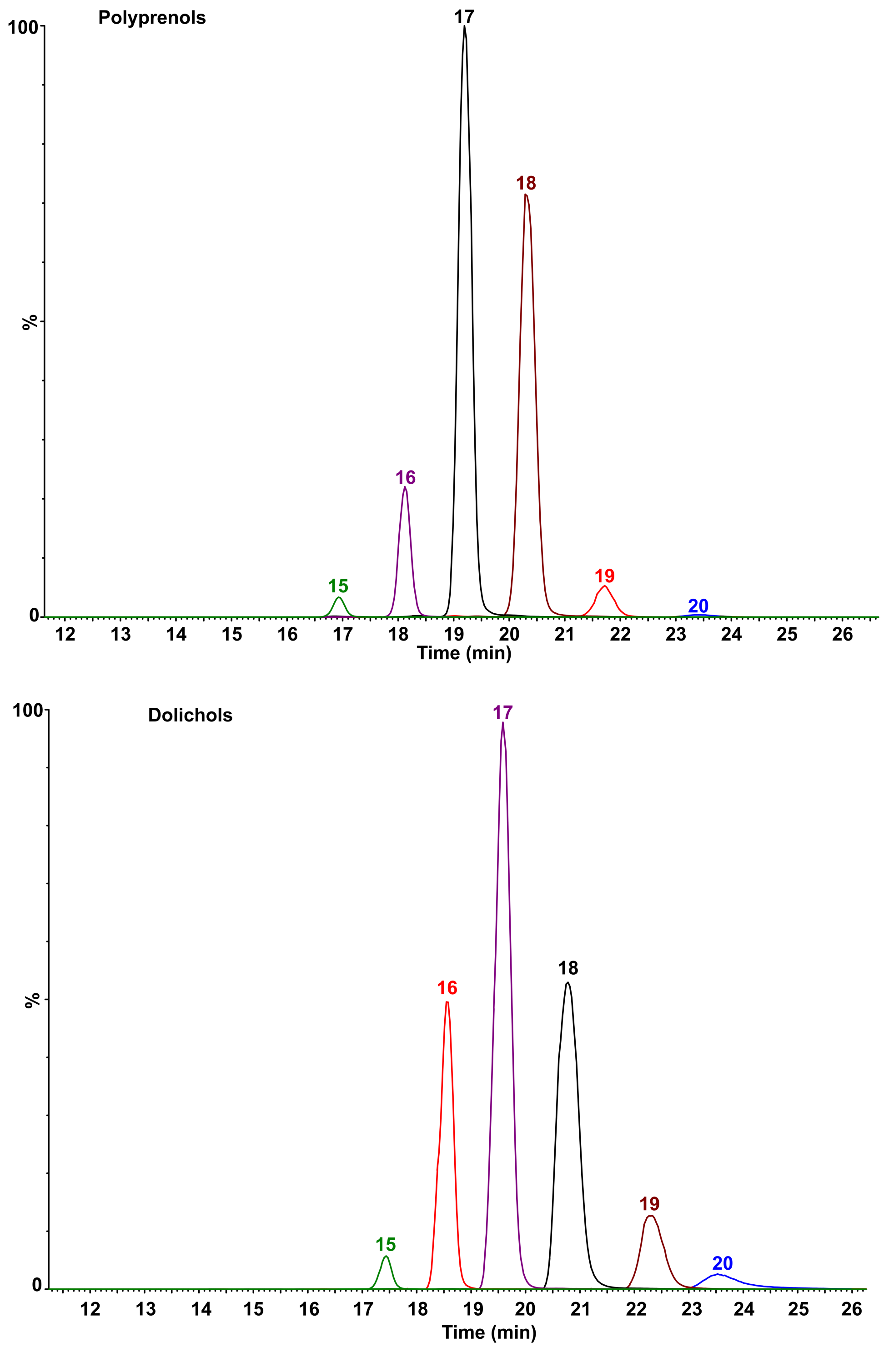
